# SSAO-mediated decrease of endothelial BDNF release affects neuronal GluA1 and PSD95 expression: a peripheral mechanism inducing NVU and CNS disturbances

**DOI:** 10.1101/2025.03.04.641410

**Authors:** Cristina Fábregas-Ordóñez, María Esteban-López, Rut Fadó, Núria Casals, Mercedes Unzeta, José Rodríguez-Alvarez, Alfredo J Miñano-Molina, Montse Solé

## Abstract

Dysfunction of the vascular system contributes to brain damage and neurodegeneration, and cerebral amyloid angiopathy (CAA) can be identified in a high percentage of Alzheimer disease (AD) brains. Blood-brain barrier (BBB) alterations can affect communication and signalling between neurovascular unit (NVU) components through changes in the release of angioneurins, among others. The two-hit vascular hypothesis of AD suggests that chronic vascular risk factors cause an early damage to the brain microvasculature (hit 1) triggering a cascade of events that leads to amyloid β-peptide (Aβ) accumulation in the brain and precipitating the Aβ-dependent pathway of neurodegeneration (hit 2). The vascular enzyme and adhesion molecule SSAO/VAP-1 plays an important role in cerebrovascular dysfunction and vascular Aβ aggregation, acting as an active player in the first of the two-hits. We generated a new NVU in vitro model to study the role of SSAO/VAP-1 on the BBB dysfunction and its effects on neurons in an AD context. We focused on the release of key angioneurins for synaptic plasticity and neuronal survival, highly compromised in AD. Our results show that SSAO/VAP-1 expression, together with the addition of Aβ_1-40_ peptide containing the Dutch mutation (used to mimic CAA condition; AβD), to cerebral endothelial cells synergistically alter the release of vascular BDNF. This alteration subsequently affects the expression of synaptic markers PSD95 and GluA1 in cortical neurons. Moreover, GluA1 and PSD95-positive neurons present reduced immunolabeling intensity of these markers when low levels of vascular BDNF are present. These results reveal vascular BDNF as a possible key element in the sensitization to Aβ due to SSAO/VAP-1 expression, impacting negatively on glutamatergic synaptic and NVU function.

**Highlights:**

- SSAO/VAP-1 expression alters BBB decreasing ZO-1, VE-Cadherin and Claudin-5 levels.
- SSAO/VAP-1 expression alters the release of vascular angioneurin BDNF.
- SSAO/VAP-1 and AβD decrease synaptic protein levels affecting glutamatergic neurons.
- Endothelial BDNF decrease induced by SSAO/VAP-1 and AβD alters glutamatergic neurons.

## 1. INTRODUCTION

The neurovascular unit (NVU) is an anatomically and functionally integrated group of different cell types that includes vascular cells (endothelial cells, vascular smooth muscle cells and perivascular pericytes), glia (astrocyte end-feet, microglia and oligodendrocytes), and neurons (Iadecola, 2010; Zlokovic, 2011). Endothelial cells within the NVU present special features, as they are not fenestrated and they present specific focal adhesions such as tight junctions or adherens junctions, resulting in a highly selective blood-brain barrier (BBB) (Abbott et al., 2010). BBB maintains the microenvironmental homeostasis in the brain, and its dysfunction contributes to brain damage and neurodegeneration (Kadry et al., 2020) through the alteration of NVU function, and an altered intercellular NVU communication and signalling by released angioneurins (Zacchigna et al., 2008; Zagrean et al., 2018).

In the aging brain and in many neurodegenerative diseases, such as Alzheimer’s disease (AD), the coexistence of cerebrovascular lesions with neurodegeneration is well documented (Govindpani et al., 2019; Kisler et al., 2017). Among AD neuropathological lesions, cerebral amyloid angiopathy (CAA) can be identified in 85-95 % of AD brains (Boyle et al., 2015) and the impact of vascular alterations on AD is greatly acknowledged at present (Iadecola, 2017). For instance, multiple pathologies with vascular components (diabetes, atherosclerosis, stroke,…) constitute risk factors for developing AD (Fillit et al., 2008; Zlokovic, 2011). The two-hit vascular hypothesis of AD suggests that chronic vascular risk factors cause an early damage to the brain microvasculature (hit 1) that can trigger a cascade of events leading to amyloid β-peptide (Aβ) accumulation in the brain and potentiating its pathological effects, precipitating the Aβ-dependent pathway of neurodegeneration (hit 2). Aβ accumulation, in turn, would strength the existing vascular and neuronal toxicity produced by hit 1, constituting a pathological feedback loop (Zlokovic, 2011). However, the molecular mechanisms underlying AD-related vascular alterations and how the NVU and brain parenchyma are affected by cerebrovascular alterations is still mostly unknown.

Vascular-adhesion protein (VAP-1) is a homodimeric glycoprotein with semicarbazide-sensitive amine oxidase enzymatic activity (SSAO, E.C 1.4.3.21), and leukocyte-adhesion function under inflammatory conditions (Jalkanen & Salmi, 2008; Smith et al., 1998). SSAO/VAP-1 is highly expressed by endothelial cells, including those in the cerebrovasculature (Castillo et al., 1998), and it is also released into blood plasma (Lyles, 1996). Its overexpression and elevated activity have been observed in cerebrovascular tissue from AD patients (Ferrer et al., 2002), and in AD patients’ blood plasma (del Mar Hernandez et al., 2005), but also in other systemic pathologies that constitute risk factors for AD, such as diabetes (Gokturk et al., 2004; Boomsma et al., 1999), atherosclerosis (Anger et al., 2007; Mészáros et al., 1999), or stroke (Hernandez-Guillamon et al., 2010; Airas et al., 2008). Interestingly, the toxicity of the products generated through its enzymatic activity are known to damage vasculature (Solé et al., 2008), to participate in Aβ aggregation (Solé et al., 2015; K. Chen et al., 2006), and are thought to contribute to pathologies progression (Unzeta et al., 2021; Pannecoeck et al., 2015). Moreover, human cerebral microvascular endothelial cells (hCMEC/D3) expressing hSSAO/VAP-1 over-release several angioneurins, such as VEGF or IL-6 among others, that activate endothelium and could be involved in disease-associated neuroinflammation (Solé et al., 2019). Altogether suggests that SSAO/VAP-1 could play an important role in cerebrovascular dysfunction and vascular Aβ aggregation, acting as an active player in the first of the two-hits of the vascular hypothesis of AD.

Endothelial-derived angioneurins can not only modulate the inflammatory state of the NVU and brain parenchyma, but also regulate neuronal maturation and function. Of particular interest in this activity is brain-derived neurotrophic factor (BDNF), due to its involvement in synaptic plasticity and neuronal survival, cell growth, and differentiation (Lynch et al., 2008). Importantly, a large part of the BDNF found in the brain is produced by cerebral endothelial cells (Monnier et al., 2017; Guo et al., 2008), and deregulation of BDNF-TrkB-associated downstream molecular pathways has been reported to be related with memory deficits, synaptic plasticity alterations and neuronal cell death observed in AD (von Bohlen und Halbach & von Bohlen und Halbach, 2018). Affectations in the BDNF-TkB signalling pathway have also been reported to modulate the levels of key synaptic proteins, such as PSD95 (Parsons et al., 2014; Yoshii & Constantine-Paton, 2014; Robinet & Pellerin, 2011; Tartaglia et al., 2001).

In the present work, we aimed to study the effect of SSAO/VAP-1-mediated endothelial damage, on neurons within the NVU. Our objective was to determine whether SSAO/VAP-1 alters BDNF release from endothelial cells and the eventual role that this alteration could have on the synaptic maturation of glutamatergic neurons in a NVU *in vitro* model, in the context of cerebrovascular pathology related to AD.

## 2. MATERIALS AND METHODS

### 2.1 Neurovascular unit *in vitro* model

To mimic a NVU environment, a new *in vitro* model was used in which the endothelial hCMEC/D3 (human cerebral microvascular endothelial cells) cell line was co-cultured with mouse mixed neuron-glia primary cells, cultured as previously described (Solé et al., 2019) with some modifications. For the NVU *in vitro* model, each individual culture was developed separately, in synchrony, until the moment to set up the co-culture as detailed as follows (Fig. S1A).

First, Mouse mixed neuron-glia primary cultures were performed as described in (Malagelada et al., 2005), with some modifications: E14.5-15.5 C57BL/6 mice (Envigo) were used; cells were seeded at 2.5 × 10^5^ cells/cm^2^ on poly-D-lysine-coated plates in Basal Medium Eagle’s (BME; Sigma) supplemented with 5% inactivated horse serum (HS; ThermoFisher Scientific), 5% FBS, 10mM glucose, and 2mM Glutamax (ThermoFisher Scientific). Cells were maintained at 37°C, in a humidified atmosphere containing 5% CO_2_ and allowed to grow until 4DIV, when half of their media was renewed by the same fresh media. At 7DIV the entire media was replaced by BME supplemented with 10% HS, 10mM glucose, 2mM Glutamax, and 10μM cytosine-β-arabinoside (Ara-C; Sigma). At 9DIV and, since then, each 3DIV, half of the media was changed by BME supplemented with 5% HS, 10mM glucose and 2mM Glutamax. At this moment, the NVU co-culture was set-up and considered as 0DIV of the co-culture (DIV-CC).

Regarding the endothelial part of the model, the original wild type (WT) hCMEC/D3 cell line was obtained from Dr. Couraud’s laboratory in Paris, France (B. Weksler et al., 2013; B. B. Weksler et al., 2005), and it does not express SSAO/VAP-1, as previously demonstrated (Sun et al., 2015). The hCMEC/D3 cell line expressing human SSAO/VAP-1 (hCMEC/D3 hSSAO/VAP-1) was generated in our laboratory as previously described (Sun et al., 2015), and it shows SSAO/VAP-1 activity levels comparable to those observed in human brain tissue (Sun et al., 2015; Hernandez-Guillamon et al., 2010). Both hCMEC/D3 cells were cultured on 150μg/mL collagen type 1 (Rat Tail, Corning)-coated plates in EBM-2 (Lonza) medium supplemented with 5% inactivated FBS (Fetal Bovine Serum, ThermoFisher Scientific), 1.4μM Hydrocortisone (Sigma), 4μg/mL Ascorbic Acid (Sigma), 1% Chemically Defined Lipid Concentrate (ThermoFisher Scientific), 1mM HEPES (ThermoFisher Scientific), 1ng/mL human Fibroblast Growth Factor-basic (bFGF; Sigma), 100U/mL penicillin (ThermoFisher Scientific), and 100μg/mL Streptomycin (ThermoFisher Scientific). In addition, cells expressing hSSAO/VAP-1 were maintained in media containing 100μg/mL geneticin (G418, ThermoFisher Scientific) to ensure plasmidic DNA maintenance. Endothelial cells were kept in an incubator at 37°C with a humidified atmosphere with 5% CO_2_. hCMEC/D3 cells were seeded at 2 × 10^5^ cells/mL and, when transwell inserts (Tw: Transwell polyethylene terephthalate transparent membrane inserts, pore size 0.4μm; Sarstedt) were used, they were additionally coated with 1% fibronectin (Sigma) in Hanks’ Balanced Salt Solution (HBSS; ThermoFisher Scientific).

For the co-culture, endothelial cells seeded on transwells were allowed to grow for 3 days in vitro (DIV). At 1 day after they reached confluence (1DAC), hCMEC/D3 cells were starved in FBS, ascorbic acid and bFGF-free EBM-2 media and put in co-culture by joining them with 12-well plates containing 9DIV mixed neuron-glia cultures.

Both parts of the system, the apical (endothelial cells) and the basolateral (neuron-glia mixed culture cells) chambers, communicate with each other through the semipermeable membrane of the Tw insert, allowing the pass of angioneurins and other molecules (Fig. S1B).

### 2.2 Cell treatments

When indicated, endothelial cells from NVU co-culture were treated with 5μM AβD (Aβ_1-40_ peptide containing the Dutch mutation, Bachem AG) at 1DIV-CC, after renewing its medium by FBS and bFGF-free conditioned EBM-2 medium. Lyophilized AβD was resuspended at 1mg/mL with 1,1,1,3,3,3-hexa-fluoro-2-propanol (HFIP, Sigma), aliquoted, evaporated and stored at −80°C until its use, being then dissolved in sterile phosphate-buffered saline (PBS), containing 0.1% ammonium hydroxide.

### 2.3 Homogenates and subcellular fractionations

To obtain complete homogenates, neuron-glia cells from NVU co-cultures were scrapped in cold NP-40 lysis buffer (20mM Tris pH 7.4, 150mM NaCl, 5mM EDTA, 1% NP-40, 1mM Na_3_VO_4_, 1mM PMSF, 1% protease inhibitor cocktail, and 1% phosphatase inhibitor cocktail). On the other hand, endothelial hCMEC/D3 cells were lysed in cold 100mM tris buffer, pH 9, supplemented with 1% protease inhibitor cocktail (Sigma). Both types of samples were then sonicated for 10 seconds. Protein extracts were quantified using Bradford reaction kit (Bio-Rad), following manufacturer’s instructions.

Post synaptic density (PSD) fraction was purified from synaptosomal membrane preparations, after a subcellular fractionation procedure with sucrose gradient. Specifically, neuron-glia mixed cells were washed with PBS and collected in HEPES buffer (4mM HEPES pH 7.4, 1% protease inhibitor cocktail (Sigma), 1% phosphatase inhibitor cocktail (Sigma), 1mM Na_3_VO_4_, 1mM NaF, 0.04% NaPPi, and 0.01% glycerol phosphate) with 0.32M sucrose (Acros), to be homogenized with a pellet pestle (Sigma) for 25 seconds. A whole homogenate aliquot (37.5μl) was taken as input sample and stored at −20°C. The rest was centrifuged for 10 minutes at 1000 x g and 4°C. Supernatant (S1) was transferred to a new tube and centrifuged for 15 minutes at 10000 x g and 4°C. Pellet (P2) was washed with HEPES-sucrose buffer and centrifuged for 15 minutes at 10000 x g and 4°C, resuspended in HEPES buffer and homogenized with the pellet pestle for 25 seconds. The homogenate was then transferred to a mini-ultracentrifuge tube (S120-AT2 rotor) and ultracentrifuged for 20 minutes at 25000 x g and 4°C. The pellet (P3) was resuspended in HEPES-sucrose buffer and loaded into a discontinuous sucrose gradient (0.8M, 1M and 1.2M) supplemented with 1% protease and phosphatase inhibitors cocktail. The gradient was ultracentrifuged in a S52-ST swinging rotor for 2 hours at 150000 x g, 4°C and lowest acceleration/brake condition. After ultracentrifugation, desired fraction (between 1M and 1.2M sucrose phases) was collected and diluted in HEPES solution to be ultracentrifuged in a S52-ST swinging rotor for 30 minutes at 150000 x g, 4°C. The pellet was resuspended in Triton-HEPES-EDTA solution (0.5% Triton X-100 (Prolabo), 50mM HEPES pH 7.4, 2 mM EDTA, 1% protease inhibitor cocktail and phosphatase inhibitor cocktail) by vortexing for 15 seconds in 2 minutes intervals during 15 minutes. This homogenate was transferred to a mini-ultracentrifuge tube (S120-AT2 rotor) and ultracentrifuged for 20 minutes at 32000 x g and 4°C. The supernatant, corresponding to the synaptic plasma membrane (SPM) fraction, was transferred to a new tube and stored at −20°C. The pellet, corresponding to the PSD fraction, was resuspended in Triton-HEPES-EDTA buffer and transferred into a new tube to be stored at −20°C. Protein concentration was determined by Pierce BCA protein assay kit (ThermoFisher Scientific).

### 2.4 Cell viability

For the lactate-dehydrogenase (LDH) activity assay, Pierce LDH Cytotoxicity Assay Kit (ThermoFisher Scientific) was used, following manufacturer’s indications. Briefly, 75 μL of media from each well was used for the assay, performed in 96-well plates, and incubated for 1 hour in the dark with LDH assay mixture. Absorbance was read at 490nm every 4 minutes (Synergy HT microplate reader; Bio-tek instruments inc), and 690nm absorbance values were subtracted as blank (KC4 analyzer software; Bio-tek instruments inc). The reaction was stopped with 75 μl/well of stop solution to take an additional end-point measurement. Cytotoxicity was inferred by the directly proportional relation to the LDH activity measured in the medium in each condition.

### 2.5 Immunoblotting

Equal amounts of protein (15 μg per line except SPM and PSD samples, with 5 μg per line), were separated in 10% SDS-PAGE and transferred onto nitrocellulose membranes (Amersham Protran 0.2 NC; Cytiva). Blots were blocked at room temperature for 1 hour with 10 % w/v non-fat dry milk, 0.1 % w/v BSA (fraction V), pH 7.4 in PBS, and incubated at 4°C overnight with primary antibody in PBS 0.1% BSA, pH7.4 after washing with 0.05 % w/v Tween-20 in PBS. Primary antibodies were: anti-GluA1 (1:1000; AB1504; RRID: AB_2113602; Merck-Millipore), anti-PSD95 (1:1000; clone 6G6-1C9; ab2723; RRID: AB_303248; Abcam), anti-β-Tubulin (1:20000; clone 5H1; 556321; RRID: AB_396360; BD Biosciences), anti-β-Actin (1:20000; clone AC-74; A2228; RRID: AB_476697; Sigma), anti-GAPDH (1:40000; clone 6C5; AM4300; RRID: AB_437392; ThermoFisher Scientific), anti-Synaptophysin (1:40000; clone SVP-38; S5768; RRID: AB_477523; Sigma), anti-Zona Occludens (ZO-1; 1:1000; 40-2200; RRID: AB_2533456; ThermoFisher Scientific), anti-VE-Cadherin (1:500; clone F-8; sc-9989; RRID: AB_2077957; Santa Cruz Biotechnology), and anti-Claudin-5 (1:500; clone A-12; sc-374221; RRID: AB_10988234; Santa Cruz Biotechnology). After washing, blots were incubated for 1 hour at room temperature with horseradish peroxidase-conjugated secondary antibodies goat anti-mouse or goat anti-rabbit (554002 and 554021; RRID: AB_395198 and RRID: 395213, respectively; BD Biosciences), diluted in blocking buffer and developed using the ECL^TM^ Western Blotting Detection Reagents (Cytiva). Protein bands were detected on X-ray films (Fuji Medical X-ray films) and semi-quantitative analysis of immunoblots was performed by densitometry using ImageJ 2.0v software (National Institutes of Health, USA; https://imageJ.nih.gov.ij/). Alternatively, Odyssey Fc Imager was used and the analysis was done through Image Studio™ Lite software 5.x (LI-COR Biosciences, USA).

### 2.6 Immunocytochemistry

Neuron-glia mixed cultures on coverslips were washed with ice-cold PBS and fixed in ice-cold 4% paraformaldehyde with 4% sucrose PBS for 15 minutes on ice. Endothelial cell cultures on coverslips were washed with ice-cold PBS and fixed in methanol-acetic acid (3:1) at −20°C for 20 minutes. Permeabilization was performed at room temperature, 20 minutes in PBS supplemented with 0.1% Triton X-100 for mixed cultures, and 30 minutes in PBS containing 0.2% Triton X-100 (PBS-T) for endothelial cells. Then, cells were washed twice with PBS prior to blocking: mixed cultures were blocked in PBS supplemented with 5% normal Goat Serum (Sigma) for 1 hour at 37°C in shaking conditions, and endothelial cells were blocked in PBS-T containing 0.2% gelatin, 20mM glycine and 5% FBS for 20 minutes. Mixed cultures were then incubated in a humidified chamber overnight at 4 °C with primary antibodies anti-GluA1 (1:1000; AB1504; RRID: AB_2113602; Merck-Millipore); anti-PSD95 (1:1000; clone 6G6-1C9; ab2723; RRID: AB_303248; Abcam); anti-Synaptophysin1 (1:1000; 101 002; RRID: AB_887905; Synaptic Systems); and anti-MAP2 (1:500, M1406; RRID: AB_477171 Sigma; and 1:1000; 188 002; RRID: AB_2138183; Synaptic Systems) diluted in PBS supplemented with 2 or 5 % normal Goat Serum (Sigma). Endothelial cells were incubated with primary antibodies anti-ZO-1 (1:100; 18-7430; RRID: AB_253048; ThermoFisher Scientific) or anti-Occludin (1:100; 33-1500; RRID: AB_2533101; ThermoFisher Scientific) in PBS-T containing 0.2% gelatin, 20mM glycine and 3% FBS overnight at 4°C with the addition of 30 minutes incubation at room temperature. After washing with PBS, cells were incubated for 1 hour (mixed cultures in humidified chamber at 37 °C and endothelial cells at room temperature) with the proper secondary antibodies (Alexa Fluor 568 Goat anti-Mouse IgG, Alexa Fluor 488 Goat anti-Rabbit IgG, and Alexa Fluor 568 Goat anti-Guinea Pig IgG; RRID: AB_144696, AB_2576217, AB_2534119 respectively; ThermoFisher Scientific) in blocking buffer. Afterwards, cells were washed twice and incubated for 5 minutes with 1μg/mL HOECHST 33258 for nuclei staining (1:10000 in PBS; H3569; ThermoFisher Scientific). Following, mixed culture coverslips were washed in PBS and mounted on glass slides with Fluoromount-G as an anti-quenching reagent (Southern Biotech). Endothelial cells coverslips were washed in PBS and mounted on slides with Mowiol mounting media (Sigma).

### 2.7 Image acquisition

Images of endothelial cells were acquired on a fluorescence microscope Nikon-ECLIPSE 90i, CFI60 infinity optical system. Neuron-glia mixed culture images were acquired on a ZEISS LSM 700 confocal laser scanning microscope using the 40X oil objective. Sequential frame acquisition was set to acquire an average of 10 planes per stack to ensure complete neurons coverage (1.0 μm interval) at 16 bits, in zoom 1, a minimum of 1024 × 1024 resolution and with laser speed value of 6. Channel gain settings were optically adjusted to minimize saturation of punctae and were maintained across experimental groups. Unmodified images were used for all analyses, which were obtained in blind manner. Fluorescent signal was analyzed from maximum intensity projections. Linear scaling was applied on images only for presentation purposes, using ImageJ software (*National Institutes of Health, USA*).

### 2.8 Image quantification and analysis

Neurons from neuron-glia mixed culture population were blindly classified depending on the mean intensity values registered for GluA1 or PSD95 labeling. For each independent experiment, GluA1-positive cells were identified and the mean intensity value from each cell was measured in the soma of each GluA1-positive cell (Fig. S2). Percentile values 25^th^ and 75^th^ were calculated from the intensity values of the control condition (non-treated co-cultures with WT endothelial cells), using GraphPad Prism 10.2.3 software (San Diego, CA, USA), and were considered the thresholds to define the subpopulation types (Fig. S2B) as low intensity (Type I; below 25^th^ percentile), intermediate intensity (Type II; between 25^th^ and 75^th^ percentile) or high intensity (Type III; higher than 75^th^ percentile). Defined thresholds were applied to the rest of experimental conditions. In case of PSD95 labeling, the same procedure was performed to define the thresholds and to classify the cells.

### 2.9 Luminex assay

Cell media from neuron-glia mixed cultures and endothelial cells, both alone or in NVU co-culture conditions, and after being treated with AβD or not, were collected and centrifuged for 10 minutes at 3000 x g and 4°C to discard possible floating death cells or debris. Supernatants were transferred to new tubes and stored at −80°C. Magnetic milliplex kit (HNDG3MAG-36K; Merck-Millipore) for BDNF detection was used following the manufacturer’s instructions. Quantification was performed with a Magpix analytical test instrument xPONENT (Luminex) and xPONENT 4.2 software (Luminex).

### 2.10 Statistical analysis

Statistical analysis was performed with Prism 10.2.3 (GraphPad Software). Data in bar graphs are reported as mean ± SEM of independent experiments. Number of independent experiments is indicated in each figure legend. Two-tailed Student’s *t*-test was used when comparing only two conditions. One-way ANOVA, followed by a Tukey’s multiple comparison *post hoc* test was used when comparing more than two conditions. A two-way ANOVA, followed by a Tukey’s multiple comparison *post hoc* test, was performed when the influence of 2 independent variables on one independent variable was analyzed. Differences were significant when p<0.05, according to the following significance levels: ****p<0.0001, ***p<0.001, **p<0.01 and *p<0.05.

## 3. RESULTS

### 3.1 Establishment and characterization of the NVU *in vitro* co-culture model

We developed a new *in vitro* NVU model based on the co-culture of the human cerebral microvascular endothelial cell line hCMEC/D3 (on the apical side) and a mouse primary neuron-glia mixed culture (on the basolateral side) (Fig. S1B), using a transwell system. To assess the development of the endothelial component according to its capacity to establish BBB properties, we monitored three tight or adherens-junction protein levels along consecutive DAC. Results indicated that hSSAO/VAP-1-expressing endothelial cells presented significantly lower basal levels of ZO-1, VE-Cadherin, and Claudin-5 at 1DAC, compared to WT cells, whose values are indicated by dotted line (Fig. 1A, B). Consequently, these SSAO/VAP-1-expressing endothelial cells were not able to establish a tight BBB, as the levels of these proteins did not increase along consecutive DAC (Fig. 1A, C). In addition, they maintained very low levels of Claudin-5, compared to each corresponding DAC in WT cells (Fig 1A, C, green bars). By contrast, the analysis along four DAC revealed that, in general, levels of junctional proteins in WT cells increased from basal levels (1DAC, represented by dotted line at 1), reaching a maximum peak around 2-3DAC (Fig. 1A, C). The lack of this effect in cells expressing hSSAO/VAP-1 indicated an alteration of BBB formation in these cells. Consistently, a two-way ANOVA statistical test revealed significant effect of genotype (hSSAO/VAP-1 expression) and DAC in this phenomenon for ZO-1 [F (1, 6) = 49.60, p < 0.0001^####^; F(3,6) = 3.367, p = 0.0448^#^], and significant genotype effect for VE-Cadherin [F (1, 32) = 23.80, p < 0.0001^####^] and Claudin-5 [F (1, 32) = 134.6, p < 0.0001^####^]. Representative immunocytochemistry images representing the structural localization of ZO-1 and Occludin (Fig. 1D) showed that only WT cells present a visible line at the cells border corresponding to tight junctions at 1, 2 and 3 DAC, while a non-organized staining was present in hSSAO/VAP-1 cells along all days. This result confirmed previously reported data showing that hSSAO/VAP-1-expressing cells present delocalization of tight-junction structure and lower transendothelial electrical resistance measurements, both necessaries for achieving a tight BBB (Solé et al., 2019). Accordingly, we postulate that hSSAO/VAP-1 expression in brain endothelial cells is associated to a defective BBB establishment, therefore fitting with a model of cerebrovascular alterations.

**Figure 1.**
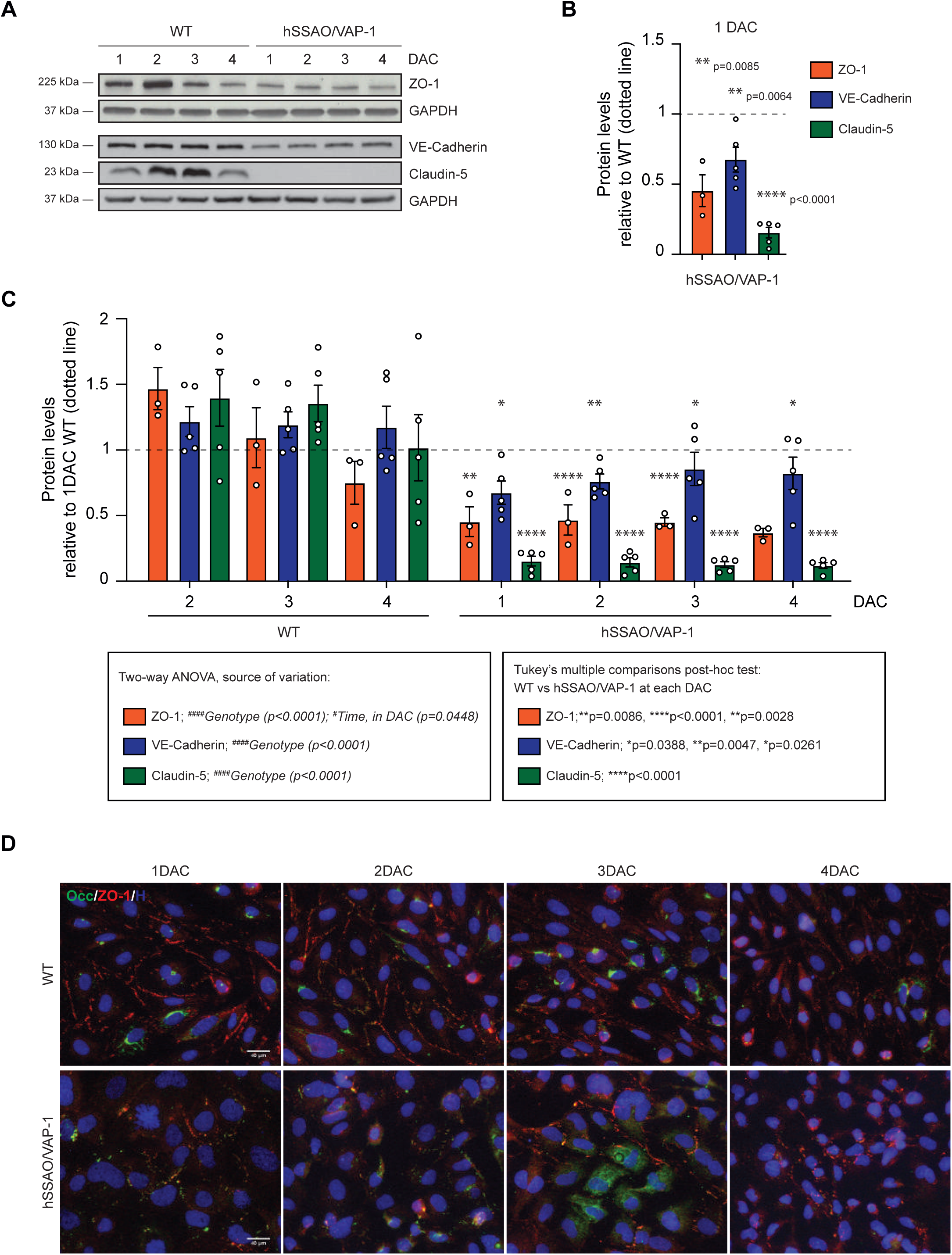
Characterization of the endothelial part of the NVU that constitutes the *in vitro* co-culture model. ***A,*** Representative immunoblots comparing tight-junction protein levels in WT and hSSAO/VAP-1-expressing endothelial cells at different DAC (days after confluence). Zonula Occludens (ZO1; ∼225-kDa band, top panel), VE-Cadherin (∼130-kDa band, middle panel) and Claudin-5 (∼23-kDa band, bottom panel. GAPDH was used as a loading control (∼36-kDa band, top and bottom panel). ***B,*** Quantification of each tight-junction protein levels at 1DAC, representing basal conditions, relative to WT (indicated by dotted line), and analyzed by Student’s t-test. ***C,*** Quantification of each tight-junction protein levels at different DACs relative to WT at 1DAC (indicated by dotted line), representing progression of BBB maturation, and analyzed by two-way ANOVA (#, source of variation) followed by Tukey’s multiple comparisons post-hoc test (*, represents WT vs hSSAO/VAP-1 at each DAC). Bars in B and C represent mean ± SEM for n=3-5 experiments. ***D,*** Representative immunofluorescence images of endothelial cells at different consecutive DAC showing Occludin (green), Zonula Occludens (ZO-1; red) and nuclei (blue) staining by Hoechst. Note the formation of organized tight-junction connections between WT endothelial cells in the ZO-1 staining in comparison to the disrupted organization of this protein in hSSAO/VAP-1-expressing endothelial cells.

On the other hand, we characterized the percentage of neurons and astrocytes in the mixed culture development and the levels of synaptic proteins at different DIV. Immunofluorescence results showed that these cultures were composed by approximately 40% of astrocytes (GFAP-positive cells) and 50% of neurons (MAP2-positive cells; Fig. 2A, B). We observed the expected increasing levels of PSD95, GluA1, and Synaptophysin during neuron development and maturation in culture (Fig. 2C-F). An inflection point in these protein levels between 9-12DIV, when they strikingly increased, indicated the beginning of the dendritic maturation process, from which high levels were maintained, denoting the achievement of a mature neuronal stage.

**Figure 2.**
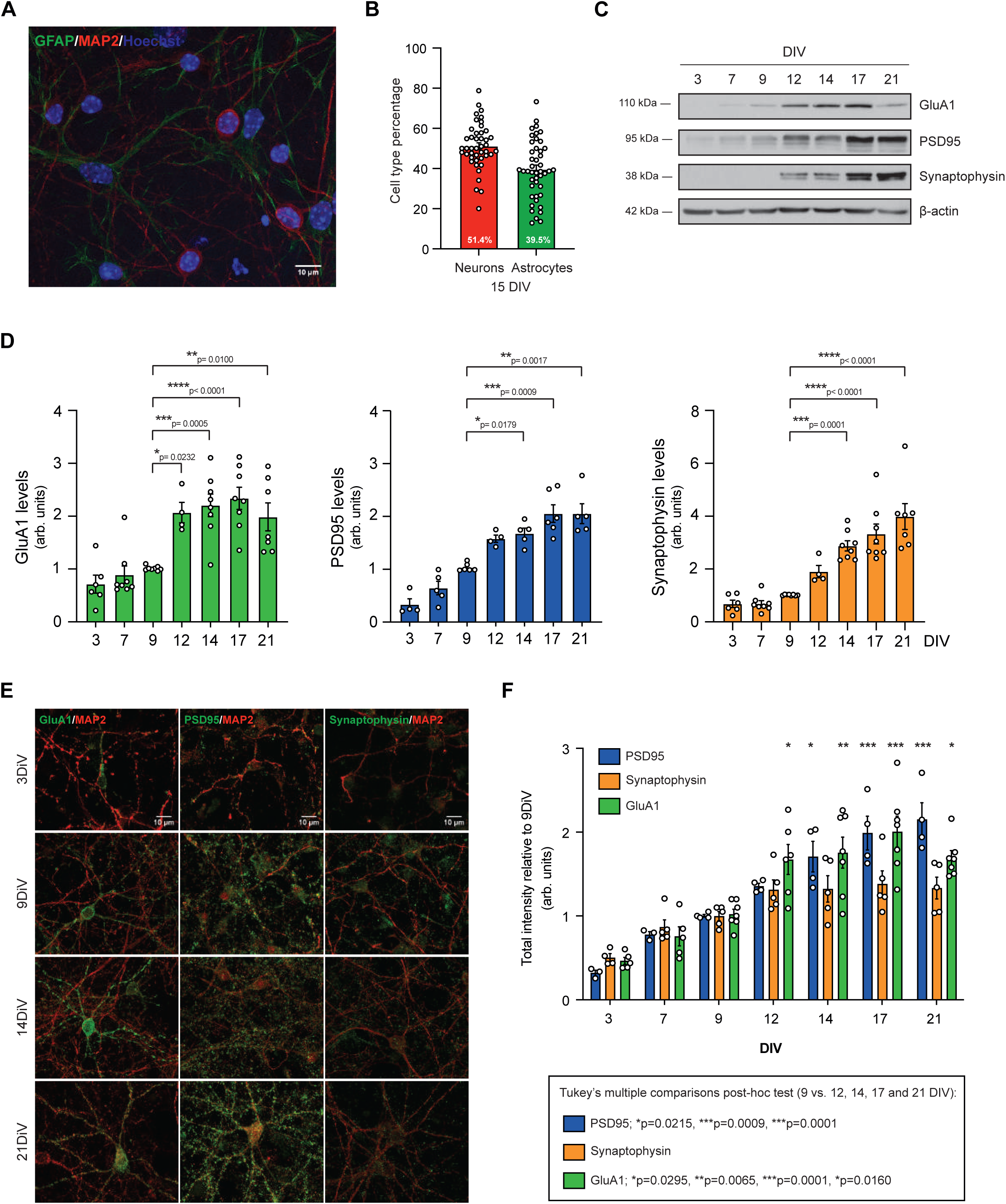
Characterization of the neuronal part of the NVU that constitutes the in vitro co-culture model. ***A,*** Representative immunofluorescence of mixed glial-neuron primary culture staining astrocytes (GFAP, green), neurons (MAP2, red) and nuclei (Hoechst, blue). ***B,*** Proportion of the of astrocyte-neuron quantification present in mixed-cortical primary cultures at 15DIV. n=45 ***C,*** Representative immunoblot images showing neuron maturity at different DIV using both pre- and post-synaptic markers GluA1 (∼110-kDa band, top panel), PSD95 (∼95-kDa band, middle-top panel), and Synaptophysin (∼38-kDa band, middle-bottom panel). β-actin was used as a loading control (∼42-kDa band, bottom panel). ***D,*** Levels of synaptic markers relative to those observed at 9DIV, corresponding to western blot quantification. Bars represent mean ± SEM; n=4-8 experiments. ***E,*** Representative confocal images of double-staining of GluA1, PSD95, and Synaptophysin with MAP2, used as a cytoskeleton marker at different DIV showing neuron maturity on mixed-cortical culture (scale bar = 10 μm). ***F,*** Quantification of each synaptic marker intensity relative to those observed at 9DIV. Bars represent mean ± SEM; n=3-7 experiments.

According to above results, we selected 1DAC for endothelial cells, and 9DIV for mixed culture as starting points to combine both cultures and establish the NVU co-culture model (Fig. S1A). Under co-culture conditions, each NVU component is able to communicate with the other by the transwell system containing a semipermeable membrane through which angioneurins can pass, simulating an *in vivo* communication between these cell types (Fig. S1B).

One of the difficulties of combining different cell types in a co-culture system is the adaptation of cell culture media, including serum concentration arrangements, as each cell type has different factor requirements to survive, especially cerebral primary cultures. An LDH cytotoxicity assay on the basolateral compartment was performed to prove mixed culture viability in medium conditions used in the co-culture, including co-culture medium (CC) with empty transwell (T.Ø), with WT endothelial cells (T.WT) or with hSSAO/VAP-1 endothelial cells seeded in transwell (T.SSAO) (Fig. S3). Results showed that neither the medium change, nor the transwell addition with endothelial cells significantly affected neuronal viability after 1 or 3 DIV of the co-culture. Compared to a cytotoxic insult (1 mM glutamate: 9.251±1.289; p<0.0001****), all the conditions released marginal levels of LDH to the medium, confirming the viability of the neuron-glia mixed cells in these co-culture conditions.

### 3.2 Endothelial cells expressing hSSAO/VAP-1 release lower levels of BDNF

BDNF is a crucial angioneurin for neuronal function, including neuronal survival and synaptic maturation and plasticity (Lynch et al., 2008). Although it has been reported that BDNF is released by endothelial cells (Monnier et al., 2017; Guo et al., 2008), it is unknown which factors can modulate BDNF release by these cells, and the impact of this source of BDNF on neuronal maturation and function.

Based on the previously observed modulation of the release of other angioneurins by hSSAO/VAP-1-expressing endothelial cells (Solé et al., 2019), we aimed to explore the eventual regulation of BDNF secretion by hSSAO/VAP-1 presence in cerebrovascular endothelial cells. To this end, we determined the amount of human (endothelial)- and mouse (mixed neuronal culture)-derived BDNF in our NVU co-culture model by Luminex assay. Results revealed that the vast majority of the BDNF present in the co-culture was released by endothelial cells (Fig. 3), as the quantification of mouse BDNF, released by the mixed culture, was not detectable (data not shown).

**Figure 3.**
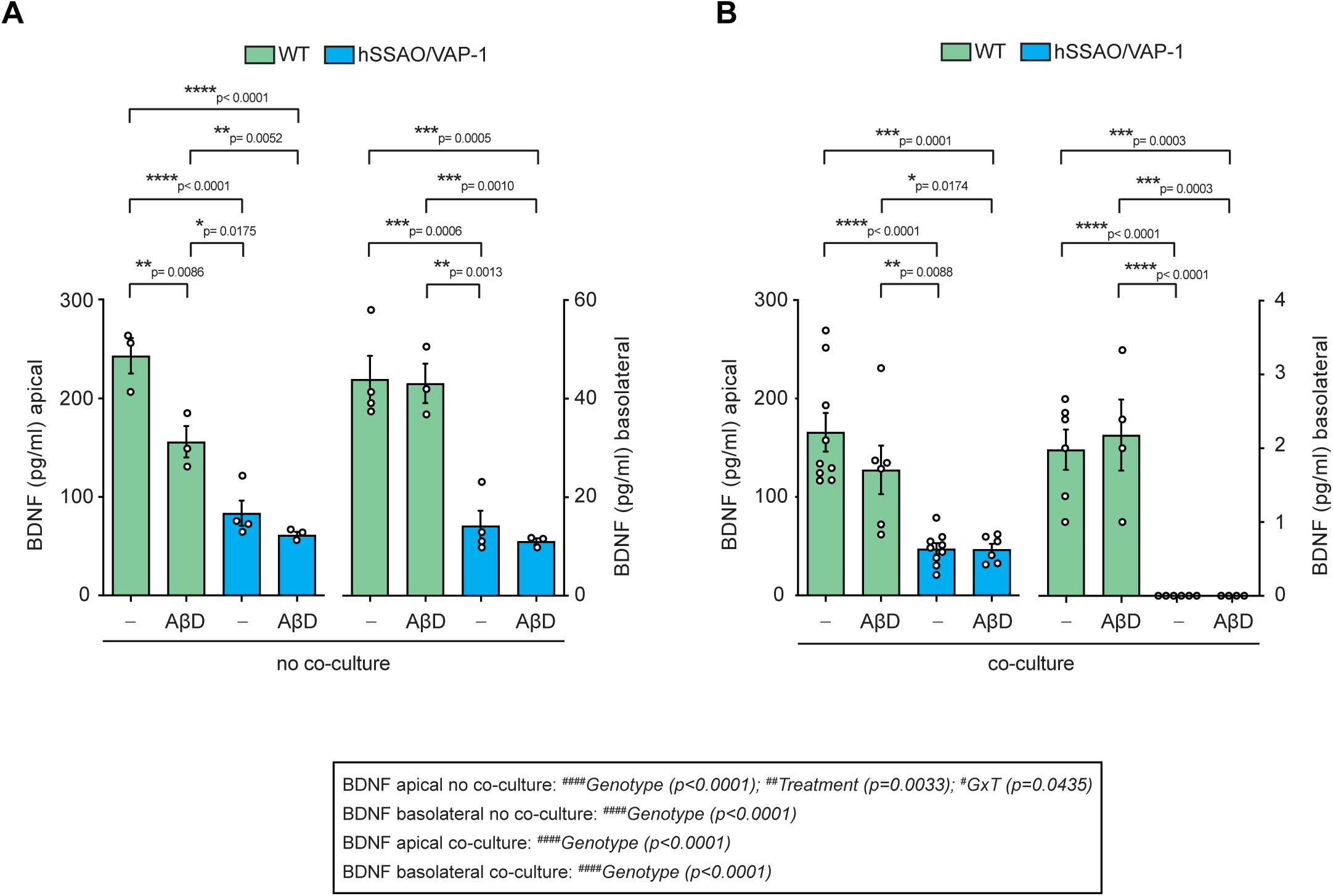
hSSAO/VAP-1 expression and AβD treatment reduce endothelial BDNF release. Levels of BDNF released into the culture medium by WT and hSSAO/VAP-1-expressing cells was quantified by Luminex in the apical and basolateral compartments. No co-culture (***A***) and co-culture (***B***) conditions were analyzed, both in basal conditions and in presence of 5μM AβD from 1 DAC, according to fig 4A. Bars represent mean ± SEM of n=3-9 independent experiments, analyzed by two-way ANOVA (#, source of variation) followed by Tukey’s multiple comparisons post-hoc test (*).

The levels of BDNF released by hSSAO/VAP-1-expressing endothelial cells in basal conditions, in both apical and basolateral compartments, were significantly lower compared to those released by WT endothelial cells, not expressing hSSAO/VAP-1, either in non-co-culture (Fig. 3A) or in co-culture conditions (Fig. 3B). In addition, levels of BDNF in basolateral compartment of hSSAO/VAP-1 NVU co-cultures were almost undetectable, suggesting a lack of this angioneurin for mixed neuronal cells in these conditions (Fig. 3B).

To evaluate changes in vascular BDNF release in the context of a cerebrovascular alteration related to AD, we exposed the endothelial part of our NVU model to the vasculotropic Dutch mutant form of Aβ (AβD) to mimic a CAA condition (Solé et al., 2015, 2019). Therefore, AβD was added to the apical (endothelial) compartment in a system not containing the neuron-glia mixed cells (no co-culture) or containing the complete NVU model (co-culture). The AβD treatment resulted in a significant reduction in the levels of BDNF detected in the apical medium of WT cells without co-culture (Fig. 3A, p=0.0086**). Similar trends were observed under co-culture conditions, showing lower BDNF amounts. The levels of BDNF released by hSSAO/VAP-1-expressing endothelial cells did not significantly change after AβD treatment, probably because they were already very low in basal conditions (Fig. 3A, B). BDNF present in basolateral compartment of the NVU model was significantly lower from that detected in the apical compartment, indicating a possible difference in the release of this angioneurin depending on endothelial cell polarity, but no differences were observed after AβD treatment.

Two-way ANOVA analysis also indicated a significant main effect of genotype in the release of BDNF in all conditions (non-co-culture apical [F(1,9)=84.36, p<0.0001^####^], basolateral [F(1,10)=68.18, p<0.0001^####^]; co-culture apical [F(1,26]=37.70, p<0.0001^####^], basolateral [F(1,16)=65.94, p<0.0001^####^]), while an additional treatment effect [F(1,9)=15.6, p=0.0033^##^] with interaction between genotype and AβD treatment [F(1,9)=5.50, p=0.0435^#^] only in the apical part of endothelial cells without co-culture.

### 3.3 Neurons co-cultured with endothelial cells expressing hSSAO/VAP-1 present decreased synaptic protein levels

We next determined the impact of the endothelial hSSAO/VAP-1 expression on synaptic protein levels in our NVU model. We analyzed the synaptic maturation of the neurons in the co-culture model by measuring the levels of glutamatergic synaptic markers according to the scheme in Fig 4A. After the apical treatment of endothelial cells with 5µM AβD for 5 days, levels of PSD95 and the GluA1 subunit of AMPA receptors were determined in total homogenates and in PSD fractions of mixed cultures in the basolateral chambers (Fig. 4B-C).

**Figure 4.**
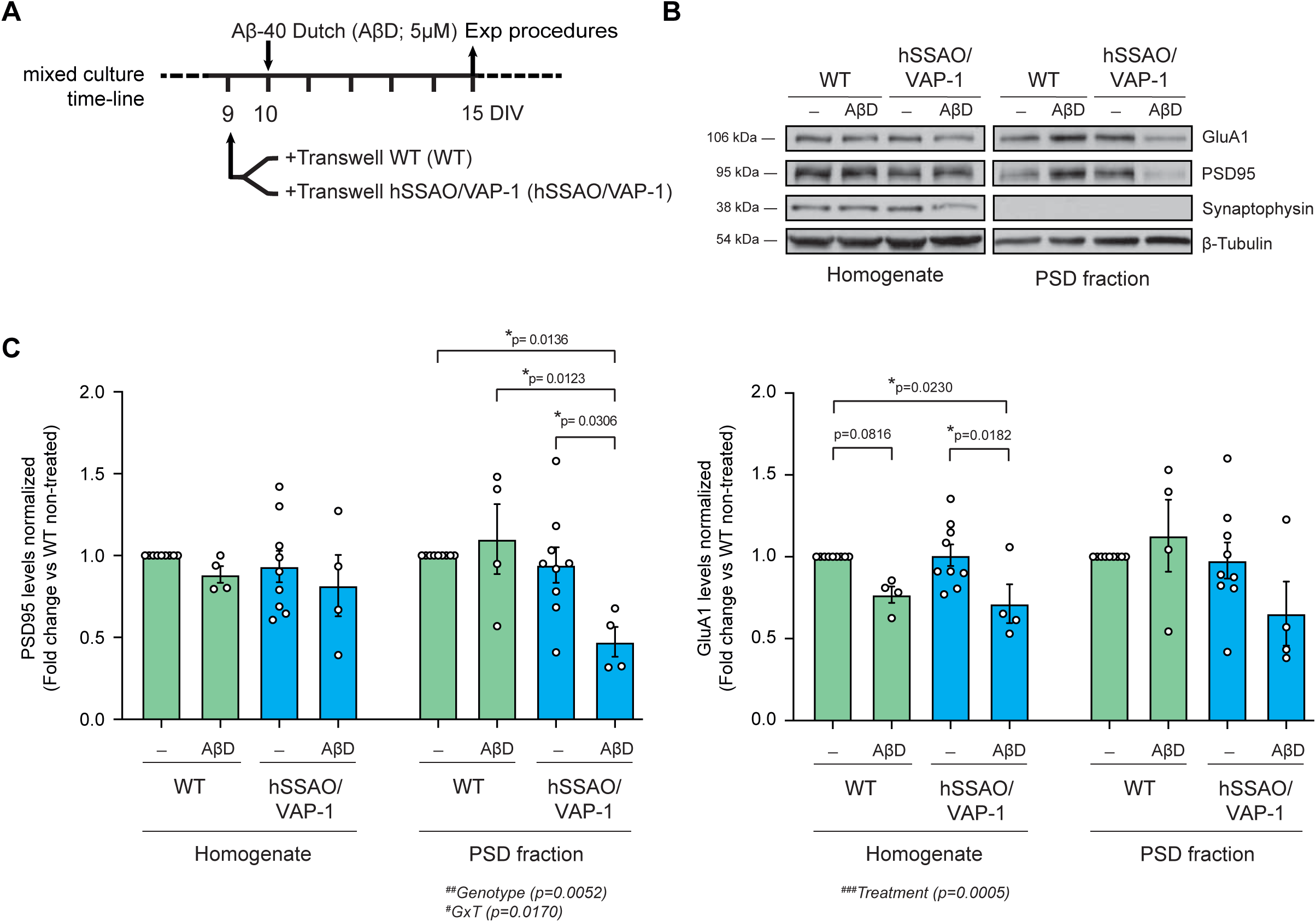
AβD treatment on endothelial cells reduces GluA1 and PSD95 protein levels in the co-cultured neurons. ***A,*** Scheme showing co-culture set up at 9DIV of neuron-glia mixed culture and 5μM AβD addition to the endothelial cells the following day in the apical compartment, with system maintenance until 15DIV, when cell lysates were collected. ***B,*** Representative immunoblot images showing levels of GluA1 (∼110-kDa band, top panel), PSD95 (∼95-kDa band, middle-top panel) and Synaptophysin (∼38-kDa band, middle-bottom panel) in total homogenates and PSD fractions from non-treated (-) and treated (AβD) endothelial cells (WT or hSSAO/VAP-1) co-cultured with neurons. Lack of synaptophysin was used as a control of PSD fraction purity. β-tubulin was used as a loading control (∼54-kDa band, bottom panel). ***C,*** Quantification of PSD95 and GluA1 protein levels relative to co-culture with non-treated WT and hSSAO/VAP-1 endothelial cells. Bars represent mean ± SEM of n=4-9 independent experiments, analyzed by two-way ANOVA (#, source of variation) followed by Tukey’s multiple comparisons post-hoc test (*).

On one hand, although no differences were observed in total homogenates, PSD95 levels (Fig. 4C, left) were significantly decreased in the PSD fraction of neurons from AβD-treated hSSAO/VAP-1 NVU co-cultures, but not in those from WT NVU co-cultures. Moreover, a two-way ANOVA analysis revealed a significant main effect of genotype (co-culture with WT or hSSAO/VAP-1) [F (1,22)=9.603, p= 0.0052^##^] and interaction between genotype and AβD treatment [F (1,22)=6.671, p=0.0170^#^], indicating that the presence of hSSAO/VAP-1 was associated with the different response to AβD treatment.

On the other hand, levels of GluA1 (Fig. 4C, right) were significantly reduced in total homogenates of neurons from hSSAO/VAP-1 NVU co-cultures treated with AβD. In this case, a two-way ANOVA analysis revealed a significant main effect of treatment [F (1,22)=16.70, p=0.0005^###^]. In the PSD fraction, GluA1 protein levels displayed similar trends to PSD95, showing a tendency to decrease in neurons from AβD-treated hSSAO/VAP-1 NVU co-cultures, compared to those from AβD-treated WT NVU co-cultures.

### 3.4 Co-cultures with hSSAO/VAP-1-expressing endothelial cells treated with AβD reduce the percentage of neurons expressing high levels of GluA1 and PSD95

To determine whether the decreased levels of PSD95 and GluA1 previously observed by western blot were a consequence of cell loss or a change in the expression levels or localization, neurons subjected to experimental co-culture conditions (Fig. 4A) were immunostained against GluA1 and PSD95 proteins. Obtained images revealed the existence of different types of GluA1-positive neurons depending on their labeling intensity. As described in the Materials and Methods, GluA1 and PSD95-positive neurons were classified according to their increasing intensity as Type I, Type II, and Type III cells (Fig. S2), being the Type I those with less intense staining and Type III those with highest intensity of staining.

The analysis of the percentage of GluA1-positive cell types showed that AβD-treated hSSAO/VAP-1 co-cultures displayed a significantly lower number of GluA1 Type III neurons compared to the basal condition (non-treated WT NVU co-cultures) (Fig. 5B, C). Similar results were observed when PSD95 Type III were analyzed (Fig. 6A and B). This effect was specific of hSSAO/VAP-1-expressing co-cultures for PSD95, while in case of GluA1, the number of Type III neurons was also reduced in WT NVU co-cultures as consequence of AβD treatment independently. Therefore, a two-way ANOVA analysis revealed a significant main effect of treatment [F (1,24)=14.99, p=0.0007^###^] and genotype [F (1,24)=7.675, p=0.0106^#^] in the changes observed in Type III neurons for GluA1 staining, and a significant main effect of genotype [F (1,16)=16.51, p=0.0009^###^] in case of PSD95. In parallel, results showed that the reduction of Type III neurons occurred with a notorious increase in the number of GluA1 and PSD95 Type I neurons, indicating an absence of cell loss but a change of status of these cells. Accordingly, a two-way ANOVA analysis revealed a significant main effect of treatment [F (1,24)=5.205, p=0.0317^#^] in case of GluA1, and a significant main effect of genotype [F (1,16)= 10.68, p=0.0048^##^] in PSD95 labeling for Type I neurons increase.

**Figure 5.**
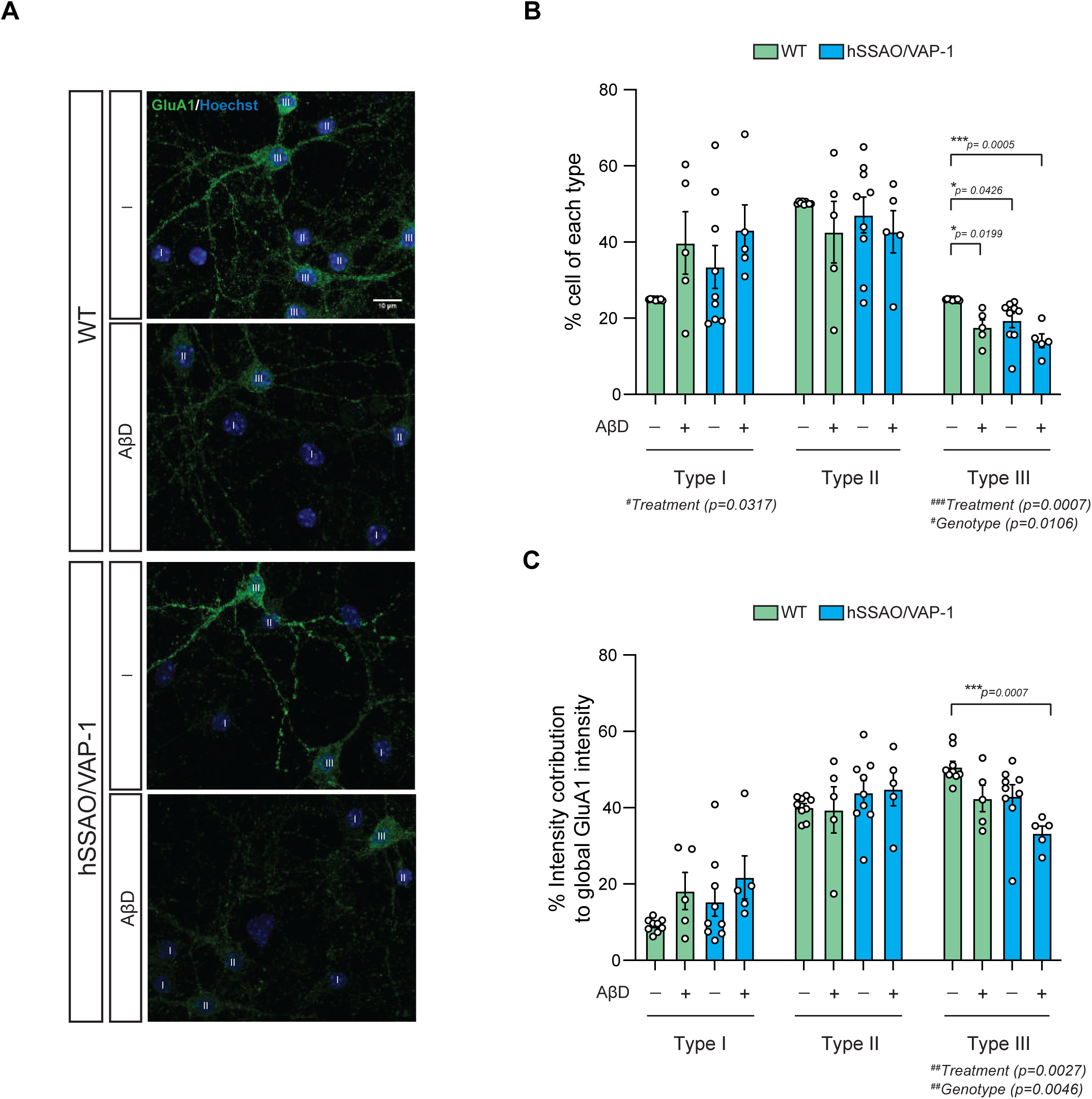
Endothelial AβD treatment reduces intensity of GluA1 staining in neurons co-cultured with hSSAO/VAP-1-expressing cells. ***A,*** Representative confocal images of neurons showing GluA1 (green) and nuclei (Hoechst, blue) staining at different experimental conditions with classification type associated to each cell body. Scale bar = 10 μm. ***B,*** Quantification of the percentage of GluA1-positive neurons classified as type I (lower) to type III (higher) intensity, according to their treatment (AβD) or not (-), and their co-culture with endothelial cells (WT or hSSAO/VAP-1). For Type III neurons: NT WT 24.88±0.05 %; AβD WT 17.61±1.96 %; NT SSAO 19.40±1.91 %; AβD SSAO 14.12±1.79 %; For Type I neurons: NT WT 24.88±0.05 %; AβD WT 39.78±8.18 %; NT SSAO 33.49±5.62 %; AβD SSAO 43.18±6.55 %. ***C,*** Quantification of the percentage of intensity contribution to global GluA1 intensity staining from non-treated (-) and treated (AβD) co-cultured neurons with endothelial cells (WT or hSSAO/VAP-1). Bars represent mean ± SEM of n=5-9 independent experiments (16 images per experiment), analyzed by two-way ANOVA (#, source of variation) followed by Tukey’s multiple comparisons post-hoc test (*). For Type III neurons: NT WT 50.72±1.43 %; AβD WT 42.42±3.46 %; NT SSAO 43.00±3.02 %; AβD SSAO 33.32±1.87 %; For Type I neurons: NT WT 9.15±0.58 %; AβD WT 18.15±4.85 %; NT SSAO 15.37±3.83 %; AβD SSAO 21.75±5.64 %.

**Figure 6.**
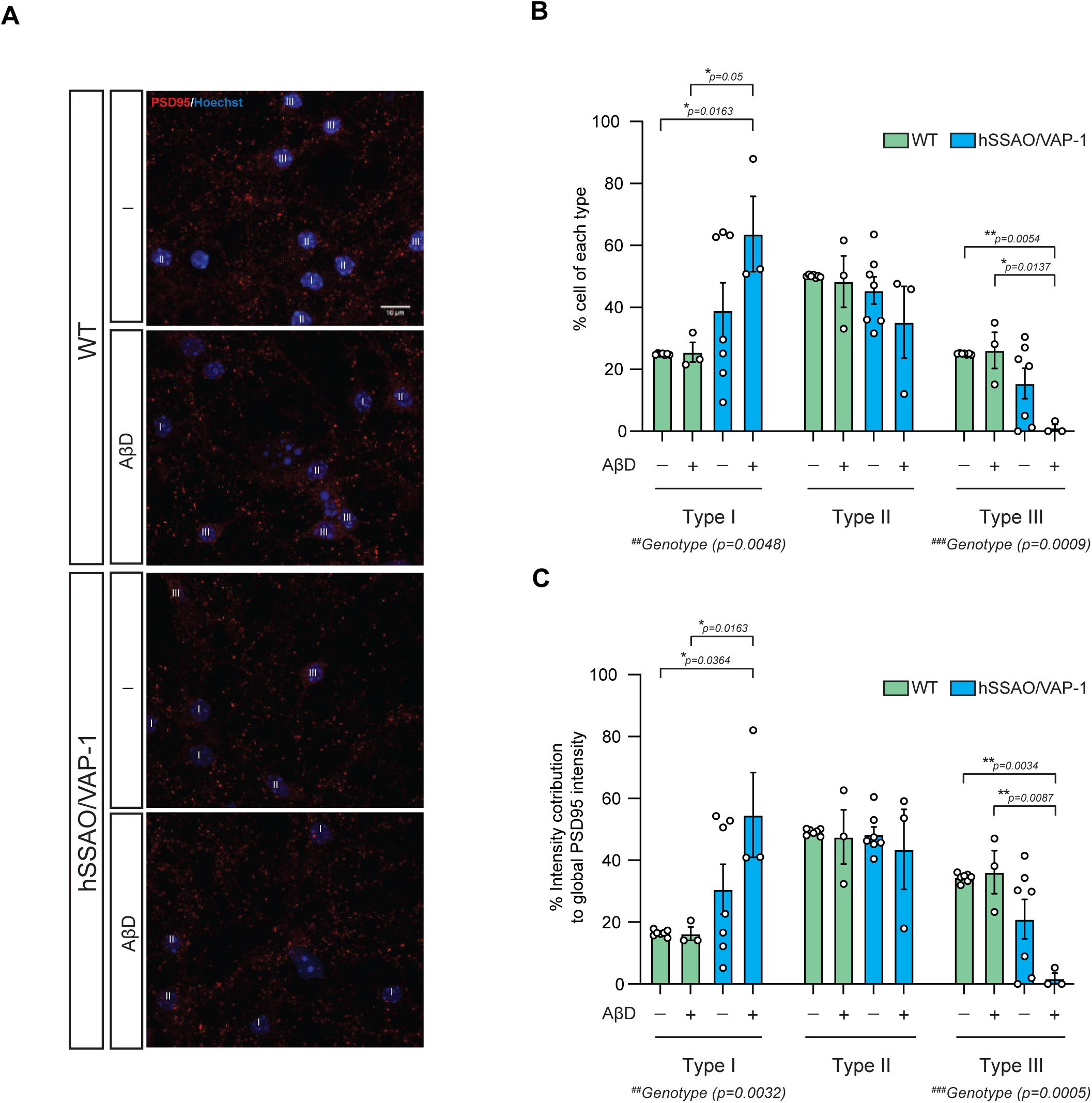
AβD treatment on endothelial cells reduces the intensity of PSD95 staining from neurons co-cultured with hSSAO/VAP-1-expressing cells. ***A,*** Representative confocal images of neurons showing PSD95 (red) and nuclei (Hoechst, blue) staining at different experimental conditions with classification type associated to each cell body. Scale bar = 10 μm. ***B,*** Quantification of the percentage of neurons for each type (low-type I to high-type III intensity) with PSD95 staining from non-treated (-) or treated (AβD) co-cultured neurons with endothelial cells (WT or hSSAO/VAP-1). For Type III neurons: NT WT 24.98±0.08 %; AβD WT 26.09±5.82 %; NT SSAO 15.45±4.88 %; AβD SSAO 1.09±1.09 %; For Type I neurons: NT WT 24.92±0.09 %; AβD WT 25.54±3.18 %; NT SSAO 39.04±8.93 %; AβD SSAO 63.71±12.14 %. ***C,*** Quantification of the percentage of intensity contribution to global PSD95 intensity staining from non-treated (-) and treated (AβD) co-cultured neurons with endothelial cells (WT or hSSAO/VAP-1). For Type III neurons: NT WT 34.42±0.52 %; AβD WT 36.15±6.94 %; NT SSAO 20.97±6.38 %; AβD SSAO 1.77±1.77 %; For Type I neurons: NT WT 16.43±0.37 %; AβD WT 16.29±2.14 %; NT SSAO 30.65±8.01 %; AβD SSAO 54.65±13.71 %. Bars represent mean ± SEM of n=3-7 independent experiments (16 images per experiment), analyzed by two-way ANOVA (#, source of variation) followed by Tukey’s multiple comparisons post-hoc test (*).

In addition, the intensity contribution of each neuronal type to the global population intensity indicates how the amount of labeling is affected in a particular type of neurons. Considering this contribution of each cell type to the intensity of the whole GluA1-positive population, results indicated that, although the number of Type III neurons in basal conditions (NT WT co-cultures) was low (20-25% of total), they contained more than 50% of the GluA1 labeling (Fig. 5BD). When co-cultured with AβD-treated hSSAO/VAP-1-expressing endothelial cells, GluA1-labeled Type III neurons contributed with significantly less intensity to the total population. A two-way ANOVA analysis revealed a significant main effect of treatment [F (1,24)=11.16, p=0.0027^##^] and genotype [F (1,24)=9.778, p=0.0046^##^]. Similar results were observed in case of PSD95 (Fig. 6AC), with a two-way ANOVA analysis revealing significant main effect of genotype [F (1,16)=18.93, p=0.0005^###^]. Consequently, type I cells intensity contribution tended to increase in co-cultures containing AβD-treated endothelial cells expressing hSSAO/VAP-1 for both GluA1 and PSD95 labeling. A two-way ANOVA analysis in case of PSD95 revealed significant main effect of genotype [F (1,16)=12.04, p=0.0032^##^].

Importantly, no differences between conditions were observed when comparing the mean intensity of each Type of neurons (Fig. S4), indicating that there is not a general reduction of intensity labeling, but an affectation of a precise group of cells.

## DISCUSSION

Angioneurins are molecular factors secreted by the integrators of the NVU to mediate the communication between the different cell types that constitute this complex structure. Angioneurins are able to simultaneously influence both vascular and neural networks during development (Adams & Eichmann, 2010), and in the adult brain (Kunze & Marti, 2019). Moreover, alteration of the cerebrovascular system, including disruption of the endothelial BBB, and alteration of angioneurin-mediated communication in the NVU, have been associated to neurodegenerative diseases such as AD and Parkinson’s disease (Noe et al., 2020; Sweeney et al., 2018). BBB integrity is essential for controlling the composition of brain interstitial fluid and therefore it is necessary, among others, for proper neuronal connectivity and synaptic function. The accumulation of Aβ around blood vessels is a hallmark of CAA, often observed in AD patients or mouse models of AD (Ma et al., 2018; Sagare et al., 2013) and can trigger NVU cells damage and dysfunction (Schultz et al., 2018). In AD, CAA induces a major disruption of BBB that contributes to cognitive decline (Greenberg et al., 2014). This BBB breakdown in AD leads to accumulation of neurotoxic material, alteration of metabolite transport and the release of pro-inflammatory angioneurins from the NVU cells (Solé et al., 2019; Sweeney et al., 2018).

The relevance of the cerebrovascular system in AD pathology is not only restricted to the established pathology, but according to the two-hit hypothesis of AD, can also contribute to its onset. This hypothesis is alternative to the classical amyloid hypothesis, only considering Aβ as initiator of the neurodegenerative process (Scheffer et al., 2021). In this regard, the two-hit hypothesis suggests that a vascular damage or alteration (hit 1) can trigger a cascade of events leading to Aβ accumulation in the brain that precipitates the Aβ-dependent pathway of neurodegeneration (hit 2), including Tau pathology and inflammation (Kisler et al., 2017). Vascular alterations include not only CAA, but also other situations with a vascular inflammatory component such as stroke, cerebral blood flow alterations, hypoperfusion, diabetes, as well as genetic and environmental factors, and they have been observed early in AD patients (Biessels, 2022; Sweeney et al., 2018)

Interestingly, disorders associated with inflammatory conditions, including AD but also other systemic pathologies, display increased plasma and/or tissular SSAO/VAP-1 levels (Danielli et al., 2022; Pannecoeck et al., 2015; Unzeta et al., 2021) that are associated to vascular damage. In a previous work, we described that the expression of SSAO/VAP-1 in endothelial cells enhances Aβ deposition on vascular cells (Solé et al., 2015), decreases tight-junction proteins (thereby producing an increase in BBB permeability), and promotes the release of several inflammatory (IL-6, IL-8, ICAM, and VCAM) or trophic factors (VEGF, TGF-β1 and NGF) (Solé et al., 2019). Therefore, SSAO/VAP-1 increased levels in the abovementioned pathologies could constitute an active player in hit 1 of the two-hit hypothesis of AD by the alterations in the endothelial angioneurins release. However, it is not known how alteration of NVU angioneurins release could affect neuronal maturation and function.

Among angioneurins, it is known that BDNF, which is involved in plasticity, neuronal survival, growth, and differentiation (Jin, 2020), is also released by endothelial cells (Marie et al., 2018; Monnier et al., 2017; Kim et al., 2004). Actually, it has been reported that a large part of the BDNF found in the brain is produced by cerebral endothelial cells (Monnier et al., 2017; Guo et al., 2008), and this was also reproduced in our NVU model. However, the role of this endothelial-derived BDNF has not been deeply studied in the NVU context. Our results, obtained in a new NVU *in vitro* model in the context of an endothelial activation due to SSAO/VAP-1 expression, revealed that hSSAO/VAP-1-expressing brain endothelial cells significantly release less BDNF than WT endothelial cells, in both conditions, cells cultured alone or in coculture, indicating that SSAO/VAP-1 expression plays a key role in BDNF release. In addition, in the NVU co-culture model, BDNF levels were always lower in the basolateral chamber when compared to the apical chamber. Whereas the lower levels observed in the basolateral medium in co-culture conditions might be explained by an uptake of the neurotrophin by the glial-neuronal culture, at present it is not known a mechanism that could explain why hSSAO/VAP-1 presence or activity might affect endothelial BDNF release. One possibility is that one of the biologically active products of its deamination activity, such as hydrogen peroxide (H_2_O_2_) or methylglyoxal, could be involved (Unzeta et al., 2021). A relationship between subtoxic levels of H_2_O_2_ and BDNF expression was reported in PC12 cells (Ogura et al., 2014). Moreover, several evidences have shown that low concentrations of H_2_O_2_ could behave as intracellular messenger with a modulatory role on different key molecules involved in signalling pathways such as NF-kB or mitogen-activated protein kinases (MAPKs) (Finkel, 1998). Further experiments need to be performed in the future to explore the role of H_2_O_2_ or other SSAO enzymatic products in the observed hSSAO/VAP-1-mediated decrease of BDNF release in endothelial cells. However, the present results together with previous ones (Solé et al., 2019) clearly show that hSSAO/VAP-1 expression in endothelial cells alters the release of angioneurins at the NVU.

BDNF and the related signalling pathways play a fundamental role in the formation, maturation, and plasticity of synapses in the central nervous system (CNS). It is well established that BDNF enhances synaptic transmission and plasticity, both by pre- and postsynaptic mechanisms (Lituma et al., 2021; Minichiello, 2009; Zakharenko et al., 2003; Levine et al., 1998; Takei et al., 1998; Vicario-Abejón et al., 1998). Moreover, BDNF is also involved in the formation and maturation of excitatory synapses in the hippocampus (Martínez et al., 1998). Most of these studies contemplated a neuronal origin for BDNF. However, although we know now that around 50% of cerebral parenchymal BDNF has an endothelial origin (Monnier et al., 2017), the contribution of endothelial BDNF to formation and plasticity of synapses is not known. In part, this was due to the difficulties to differentially identify endothelial BDNF from other cellular sources in the NVU (neurons and glial cells).

Here, we were able to avoid this caveat by using a NVU experimental model with human endothelial cells and a glia-neurons co-culture from mouse, and specifically quantify human BDNF. This experimental approach allowed to test whether the reduction in endothelial BDNF released by SSAO/VAP-1-expressing vascular cells had an impact on excitatory synapses. For that purpose we decided to monitor the postsynaptic scaffolding proteins PSD95 and GluA1 (a subunit of the glutamatergic AMPA receptors) as post-synaptic markers (Chen et al., 2005) in our NVU model.

Our data indicates that NVU co-cultures containing hSSAO/VAP-1-expressing human brain endothelial cells, showing a decreased BDNF release, produced a marked reduction in PSD95 protein levels in the PSD fraction. Our result is consistent with the reported modulation of PSD95 by BDNF in visual cortical neurons (Yoshii & Constantine-Paton, 2014) or hippocampal cultured neurons (Hu et al., 2011). This reduction was specifically observed in co-cultures containing vasculotropic AβD-treated hSSAO/VAP-1-expressing endothelial cells (to mimic a CAA condition), suggesting that the presence of hSSAO/VAP-1 could sensitize neurons to this aversive stimulus. Overall, the data is in consonance with the two-hit vascular hypothesis of AD that claims that Aβ-independent (hit 1) and Aβ-dependent (hit 2) mechanisms interrelate and merge on blood vessels, leading synergistically and/or independently to neuronal and synaptic dysfunction (Kisler et al., 2017).

When analyzing the nature of this synaptic protein decrease, complementary studies by immunocytochemistry and confocal microscopy allowed us to identify different types of neurons based on its levels of staining for PSD95 or GluA1 (Type I, II and III; from lower to higher levels of expression). Thus, our results showed that the lower release of BDNF by the AβD-treated hSSAO/VAP-1 cells was associated to a delay in excitatory neuronal maturation that was evidenced by a decrease in Type III neurons (with highest immunoreactivity to GluA1 or PSD95) and an increase in Type I neurons (with the lowest immunoreactivity to PSD95 or GluA1). This result suggests, as previously mentioned, that both AβD treatment and hSSAO/VAP-1 expression-mediated reduction of BDNF release from endothelial cells might be involved in the delay of the *in vitro* synaptic maturation of glutamatergic neurons, supporting the important role that endothelial BDNF has on synaptic function, at least, in glutamatergic neurons. Although it is beyond our study, it is interesting to point out that the reduction of endothelial BDNF release at the NVU would impact neuronal function in several ways. It has been already reported, by using conditioned media, that endothelial BDNF protects cultured neurons from insults like ischemia-hypoxia and amyloid neurotoxicity (Guo et al., 2008). Moreover, as previously indicated, BDNF has a preeminent role in the formation and maturation of synapses (Zagrebelsky & Korte, 2014; Martínez et al., 1998) and synaptic plasticity (Minichiello, 2009). Indeed, low levels of BDNF reduced spine density, that is associated to an increase in spine length and a decrease of spine head width, and BDNF is required for activity-dependent maintenance of spines in mature neurons (Kellner et al., 2014; Rauskolb et al., 2010). Reduction of BDNF levels also results in decrease in synaptic AMPA receptors and an impairment of hippocampal synaptic plasticity (Català-Solsona et al., 2023; Novkovic et al., 2015; G. Chen et al., 1999). These effects on synaptic stability and plasticity would be of particular importance in the AD brain where: a) oligomeric forms of Aβ are known to alter synaptic function and to promote spine loss (Shankar et al., 2008; Hsieh et al., 2006) and b) an increase in SSAO/VAP-1 has been observed in the neurovascular system in patients with AD (Ferrer et al., 2002), and enhanced in AD patients with diabetes (Valente et al., 2012). In AD transgenic mice models, BDNF levels are reduced in early stages of the disease (España et al., 2010). In healthy aged humans, it was reported a negative correlation between serum BDNF levels and age (Ziegenhorn et al., 2007). AD patients present diminished serum, plasma and cerebrospinal fluid BDNF concentrations compared to controls (Gezen-Ak et al., 2013). Thus, it seems reasonable that the observed reduction of BDNF would have a major contribution in the dysfunction of glutamatergic synapses in early AD prior to the neurodegenerative stage. As our results suggest, these studies point to the important role of endothelial BDNF for the synapse maintenance and hippocampal synaptic plasticity, although we are aware that an important limitation of our study is the lack of a cause-effect response prove of the absence of this endothelial BDNF in the alterations observed in the mixed neuronal culture of our NVU model.

Altogether, our data suggest that the presence of AβD-treated hSSAO/VAP-1-expressing endothelial cells in NVU co-cultures elicits a decrease in the abundance of these synaptic markers. In addition, the observed balance between Type III and Type I neurons suggests that this loss of synaptic markers is not associated to a neuronal cell death but to a delay in the formation and maturation of glutamatergic neurons in this new *in vitro* model of NVU. Moreover, the fact that in some cases differences are mainly found when both hSSAO/VAP-1 and AβD are present let us hypothesize that hSSAO/VAP-1 could act as a sensitizing effector in co-cultured neurons to an aversive stimulus such as AβD presence. Thus, these data support that endothelial BDNF is important in the formation and maturation of glutamatergic synapses and that the increase in SSAO/VAP-1 expression reported in the brain vessels of AD patients could have, among others, a negative impact in glutamatergic synaptic function.

Considering that SSAO/VAP-1 is increased in multiple disorders with an inflammatory component, and in vascular risk factors for AD, along with the already reported bidirectional relationship between SSAO/VAP-1 increase and Aβ aggregation, we could speculate the scenario shown in figure 7. We propose that under a persistent inflammatory state or in chronical diseases considered vascular risk factors for AD, SSAO/VAP-1 would be overexpressed, leading to vascular damage. This vascular damage would cause dysfunction of the BBB, an additional overexpression of SSAO/VAP-1, an alteration in the pattern of angioneurins release, and the aggregation of vascular Aβ, with consequences for the NVU and neurons dysfunction. Thus, our results agree with previous claims (Govindpani et al., 2019) about the possibility to achieve great short and long term improvement outcomes by treating vascular dysfunction and inflammation in early stages of AD, or even in pre-symptomatic situations.

**Figure 7.**
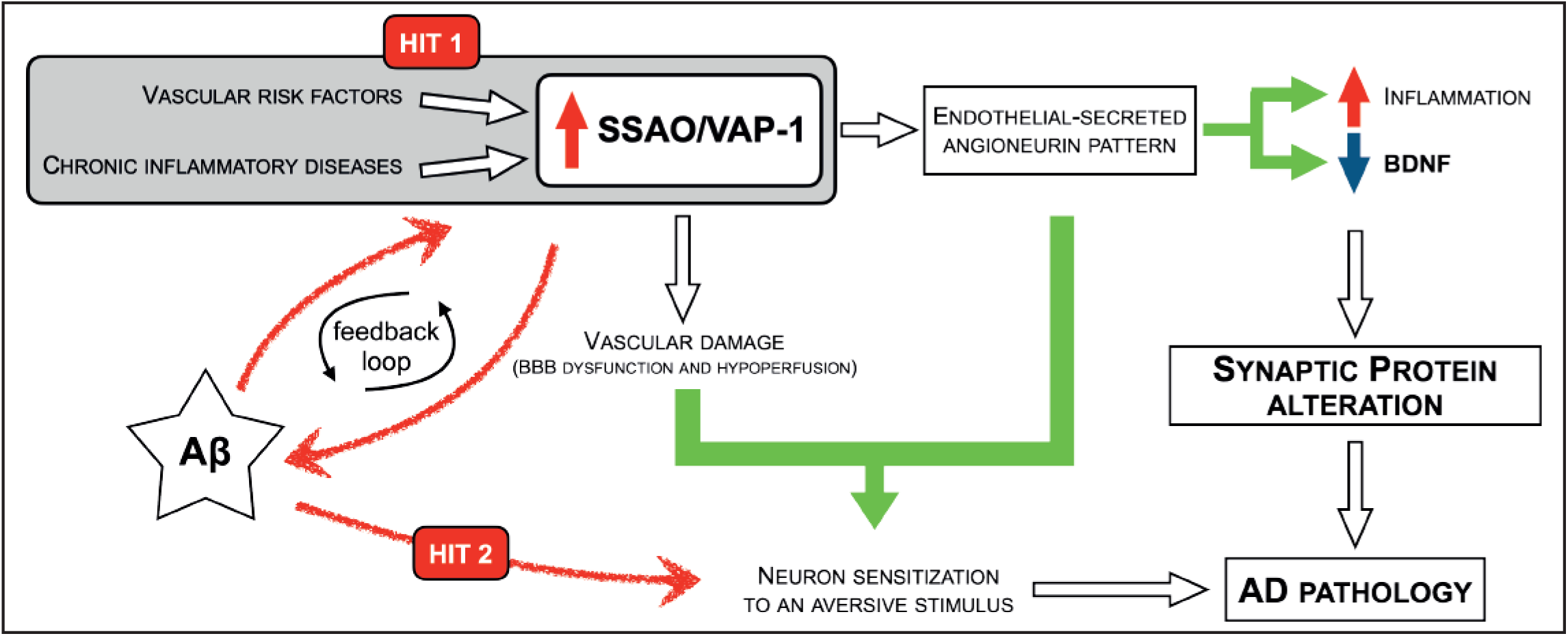
Summary diagram. Persistent inflammatory state or chronical diseases, (considered vascular risk factors for AD) will promote vascular damage by SSAO/VAP-1 overexpression (HIT 1). This vascular damage causes dysfunction of the BBB. Moreover, SSAO/VAP-1 would potentiate the aggregation of Aβ (HIT 2), which in turn promotes SSAO/VAP-1 overexpression (HIT 1), generating a feed-back loop that alters angioneurins release pattern mediating alteration of excitatory synaptic proteins. All these effects could increase neuron sensitization to an aversive stimulus contributing to AD pathology.

## Acknowledgments

This work was partially supported by grants from Fundació La Marató de TV3 (2014/3610), MCIN/AEI/10.13039/501100011033 and by “ERDF A way of making Europe” by the European Union (SAF2017-89271-R and PID2020-11751ORB-100 to J.R.A), CIBERNED (CB06/05/0042 to J.R.A) and CIBEROBN (CB06/03/0001 to N.C). C.F was supported by a PIF fellowship from Departament de Bioquímica i Biologia Molecular, in UAB.

---

**Supplementary figure 1.**
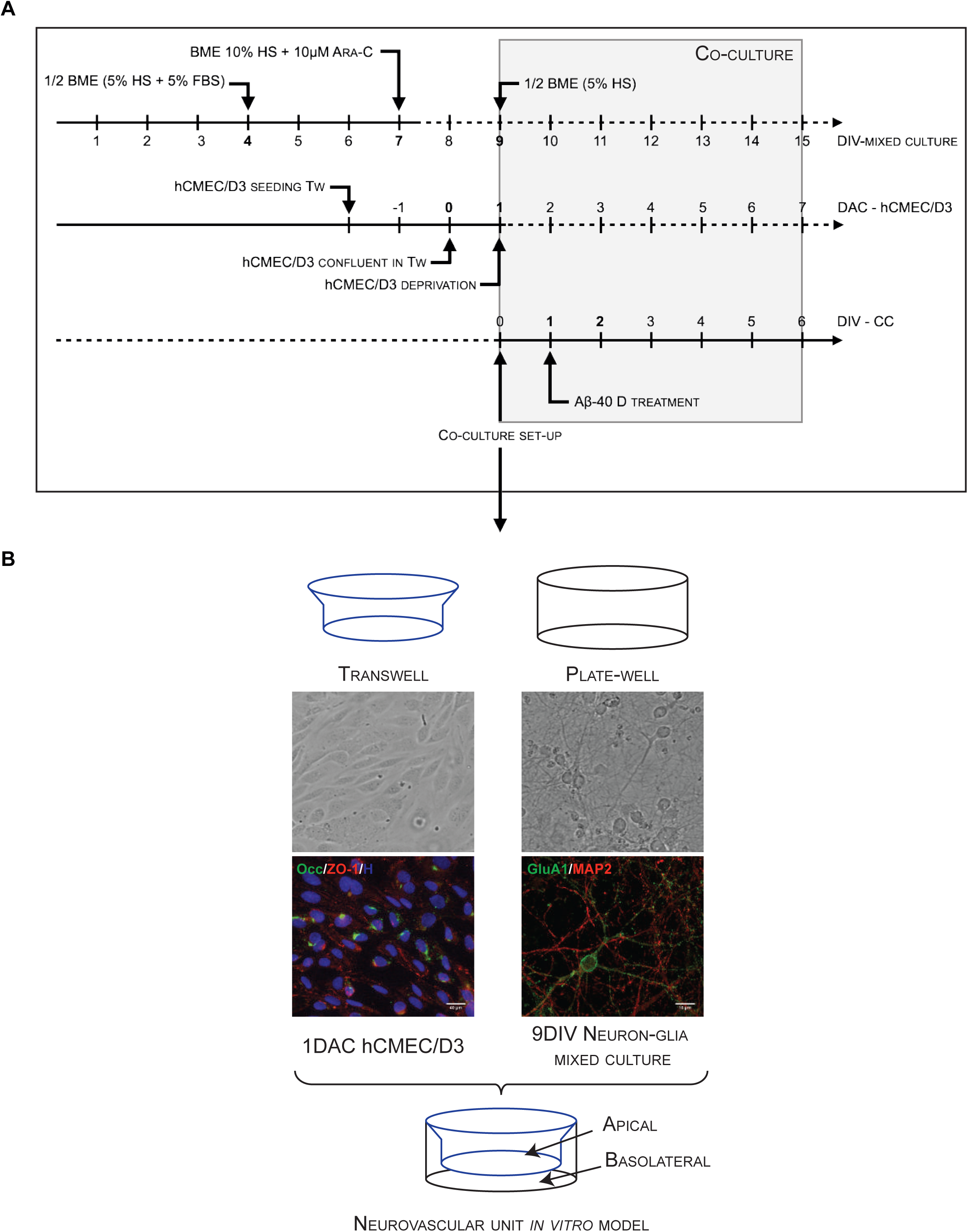
Neurovascular unit (NVU) *in vitro* model ***A,*** Extended experimental timeline. Upper arrow represents neuron-glia mixed culture timeline, indicated as days in vitro (DIV). Medium changes needed at each time point are indicated. Middle arrow represents hCMEC/D3 endothelial cell line culture timeline. Seeding, confluence, and deprivation steps are indicated and were synchronized to mixed culture development. Day 0 is considered the day when cells achieve confluence and after this, time of this culture is indicated as days after confluence (DAC). Lower arrow represents the timeline once co-culture system is set up, and time is indicated as days in vitro since co-culture (DIV - CC). Treatments are indicated the day they were performed after setting up the co-culture. ***B,*** Co-culture experimental scheme with representative phase contrast and immunocytochemistry images from WT endothelial cells (Occ: Occludin, ZO-1: Zonula occludens, H: Hoechst) and neuron-glia mixed culture (GluA1, MAP2) before setting up the co-culture. According to the time line in **A**, WT endothelial cells seeded in the transwell device (TW) were placed inside the neuron-glia mixed culture well. Both parts can communicate through the semipermeable membrane at the bottom of the transwell.

**Supplementary Figure 2.**
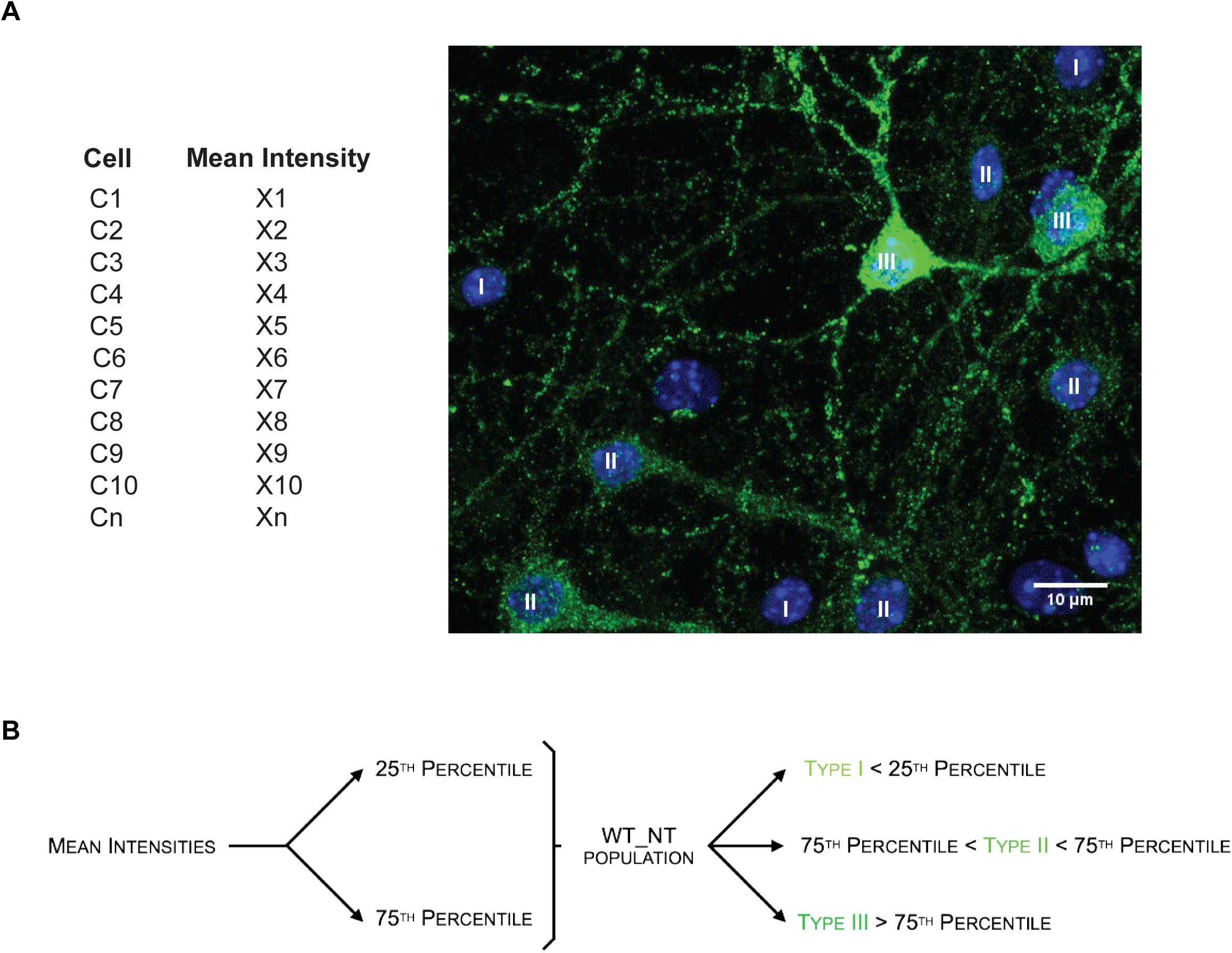
Classification of subpopulation types according to GluA1-labeling mean intensity values. ***A,*** Table showing theoretical GluA1-positive cells (C1, C2, Cn) and the mean intensity of each one (X1, X2, Xn), with representative image showing the examples of the three subtypes. Mixed primary cells co-cultured with non-treated WT endothelial cells were used as control condition. Scale bar = 10μm. ***B***, Mean intensity values from control condition were used to establish the thresholds (25^th^ and 75^th^ percentile intensity values) and to generate the subpopulation types (Type I, Type II, Type III) for each independent culture. These threshold values were applied to the rest of conditions of the same experiment to classify cells into the proper subpopulation types.

**Supplementary Figure 3.**
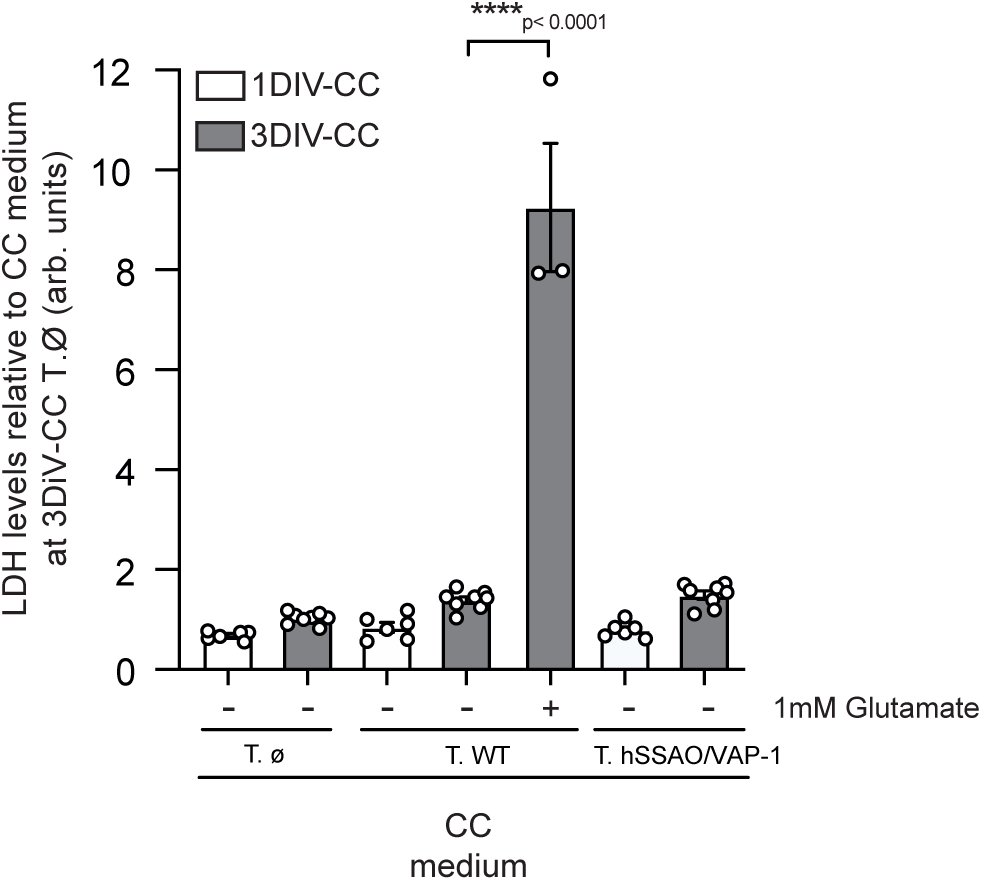
LDH levels released by mixed culture. Co-culture medium (CC) was tested and compared with an excitotoxic insult. Bars represent mean ± SEM for n=3-8 independent experiments. Data was analyzed by one-way ANOVA analysis followed by Tukey post-hoc test.

**Supplementary figure 4.**
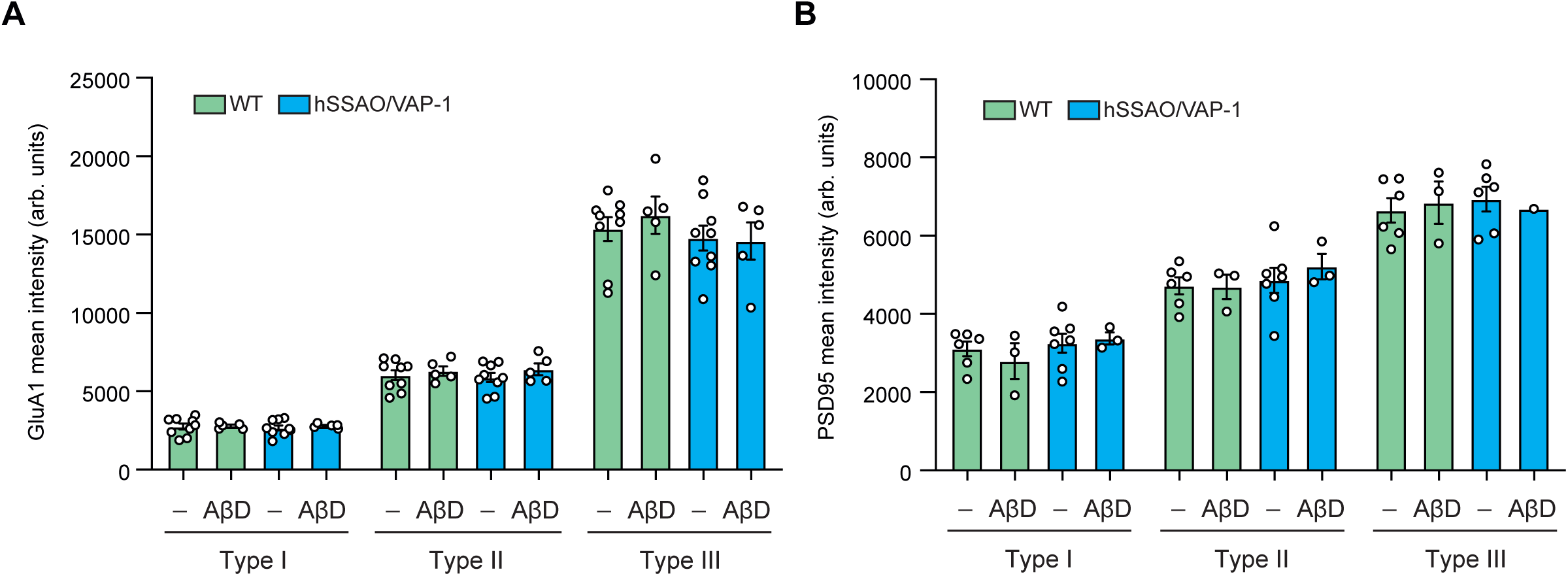
Global mean intensity of GluA1 (***A***) and PSD95 (***B***) for each subpopulation type. Bars represent mean ± SEM for n=3-9 independent experiments. Data was analyzed by two-way ANOVA analysis followed by Tukey post-hoc test.

## REFERENCES

Abbott, N. J., Patabendige, A. A. K., Dolman, D. E. M., Yusof, S. R., & Begley, D. J. (2010). Structure and function of the blood–brain barrier. Neurobiology of Disease, 37(1), 13–25. 10.1016/j.nbd.2009.07.030

Adams, R. H., & Eichmann, A. (2010). Axon guidance molecules in vascular patterning. Cold Spring Harbor perspectives in biology, 2(5). 10.1101/CSHPERSPECT.A001875

Airas, L., Lindsberg, P. J., Karjalainen-Lindsberg, M.-L., Mononen, I., Kotisaari, K., Smith, D. J., & Jalkanen, S. (2008). Vascular adhesion protein-1 in human ischaemic stroke. Neuropathology and Applied Neurobiology, 34(4), 394–402. 10.1111/j.1365-2990.2007.00911.x

Anger, T., Pohle, F. K., Kandler, L., Barthel, T., Ensminger, S. M., Fischlein, T., Weyand, M., Stumpf, C., Daniel, W. G., & Garlichs, C. D. (2007). VAP-1, Eotaxin3 and MIG as potential atherosclerotic triggers of severe calcified and stenotic human aortic valves: Effects of statins. Experimental and Molecular Pathology, 83(3), 435–442. 10.1016/j.yexmp.2007.02.008

Biessels, G. J. (2022). Alzheimer’s disease, cerebrovascular disease and dementia: Lump, split or integrate? Brain, 145(8), 2632–2634. 10.1093/brain/awac228

Boomsma, F., van den Meiracker, A. H., Winkel, S., Aanstoot, H. J., Batstra, M. R., Man in ’t Veld, A. J., & Bruining, G. J. (1999). Circulating semicarbazide-sensitive amine oxidase is raised both in Type I (insulin-dependent), in Type II (non-insulin-dependent) diabetes mellitus and even in childhood Type I diabetes at first clinical diagnosis. Diabetologia, 42(2), 233–237. 10.1007/s001250051143

Boyle, P. A., Yu, L., Nag, S., Leurgans, S., Wilson, R. S., Bennett, D. A., & Schneider, J. A. (2015). Cerebral amyloid angiopathy and cognitive outcomes in community-based older persons. Neurology, 85(22), 1930–1936. 10.1212/WNL.0000000000002175

Castillo, V., Lizcano, J. M., Visa, J., & Unzeta, M. (1998). Semicarbazide-sensitive amine oxidase (SSAO) from human and bovine cerebrovascular tissues: Biochemical and immunohistological characterization. Neurochemistry International, 33(5), 415–423. 10.1016/S0197-0186(98)00045-X

Català-Solsona, J., Lituma, P. J., Lutzu, S., Siedlecki-Wullich, D., Fábregas-Ordoñez, C., Miñano-Molina, A. J., Saura, C. A., Castillo, P. E., & Rodriguez-Álvarez, J. (2023). Activity-Dependent Nr4a2 Induction Modulates Synaptic Expression of AMPA Receptors and Plasticity via a Ca2+/CRTC1/CREB Pathway. The Journal of Neuroscience: The Official Journal of the Society for Neuroscience, 43(17), 3028–3041. 10.1523/JNEUROSCI.1341-22.2023

Chen, G., Kolbeck, R., Barde, Y. A., Bonhoeffer, T., & Kossel, A. (1999). Relative contribution of endogenous neurotrophins in hippocampal long-term potentiation. The Journal of Neuroscience: The Official Journal of the Society for Neuroscience, 19(18), 7983–7990. 10.1523/JNEUROSCI.19-18-07983.1999

Chen, K., Maley, J., & Yu, P. H. (2006). Potential inplications of endogenous aldehydes in beta-amyloid misfolding, oligomerization and fibrillogenesis. Journal of Neurochemistry, 99(5), 1413–1424. 10.1111/j.1471-4159.2006.04181.x

Chen, X., Vinade, L., Leapman, R. D., Petersen, J. D., Nakagawa, T., Phillips, T. M., Sheng, M., & Reese, T. S. (2005). Mass of the postsynaptic density and enumeration of three key molecules. Proceedings of the National Academy of Sciences of the United States of America, 102(32), 11551–11556. 10.1073/pnas.0505359102

Danielli, M., Thomas, R. C., Quinn, L. M., & Tan, B. K. (2022). Vascular adhesion protein-1 (VAP-1) in vascular inflammatory diseases. Vasa, 51(6), 341–350. 10.1024/0301-1526/a001031

del Mar Hernandez, M., Esteban, M., Szabo, P., Boada, M., & Unzeta, M. (2005). Human plasma semicarbazide sensitive amine oxidase (SSAO), β-amyloid protein and aging. Neuroscience Letters, 384(1), 183–187. 10.1016/j.neulet.2005.04.074

España, J., Valero, J., Miñano-Molina, A. J., Masgrau, R., Martín, E., Guardia-Laguarta, C., Lleó, A., Giménez-Llort, L., Rodríguez-Alvarez, J., & Saura, C. A. (2010). Beta-Amyloid disrupts activity-dependent gene transcription required for memory through the CREB coactivator CRTC1. J Neurosci, 30(28), 9402–9410. 10.1523/JNEUROSCI.2154-10.2010

Ferrer, I., Lizcano, J. M., Hernández, M., & Unzeta, M. (2002). Overexpression of semicarbazide sensitive amine oxidase in the cerebral blood vessels in patients with Alzheimer’s disease and cerebral autosomal dominant arteriopathy with subcortical infarcts and leukoencephalopathy. Neuroscience Letters, 321(1), 21–24. 10.1016/S0304-3940(01)02465-X

Fillit, H., Nash, D. T., Rundek, T., & Zuckerman, A. (2008). Cardiovascular risk factors and dementia. The American Journal of Geriatric Pharmacotherapy, 6(2), 100–118. 10.1016/j.amjopharm.2008.06.004

Finkel, T. (1998). Oxygen radicals and signaling. Current Opinion in Cell Biology, 10(2), 248–253. 10.1016/S0955-0674(98)80147-6

Gezen-Ak, D., Dursun, E., Hanaǧasi, H., Bilgiç, B., Lohman, E., Araz, Ö. S., Atasoy, I. L., Alaylioǧlu, M., Önal, B., Gürvit, H., & Yilmazer, S. (2013). BDNF, TNFα, HSP90, CFH, and IL-10 serum levels in patients with early or late onset Alzheimer’s disease or mild cognitive impairment. Journal of Alzheimer’s disease: JAD, 37(1), 185–195. 10.3233/JAD-130497

Gokturk, C., Nordquist, J., Sugimoto, H., Forsberg-Nilsson, K., Nilsson, J., & Oreland, L. (2004). Semicarbazide-sensitive amine oxidase in transgenic mice with diabetes. Biochemical and Biophysical Research Communications, 325(3), 1013–1020. 10.1016/j.bbrc.2004.10.140

Govindpani, K., McNamara, L. G., Smith, N. R., Vinnakota, C., Waldvogel, H. J., Faull, R. L., & Kwakowsky, A. (2019). Vascular Dysfunction in Alzheimer’s Disease: A Prelude to the Pathological Process or a Consequence of It? Journal of Clinical Medicine, 8(5), Article 5. 10.3390/jcm8050651

Greenberg, S. M., Salman, R. A.-S., Biessels, G. J., Buchem, M. van, Cordonnier, C., Lee, J.-M., Montaner, J., Schneider, J. A., Smith, E. E., Vernooij, M., & Werring, D. J. (2014). Outcome markers for clinical trials in cerebral amyloid angiopathy. The Lancet Neurology, 13(4), 419–428. 10.1016/S1474-4422(14)70003-1

Guo, S., Kim, W. J., Lok, J., Lee, S. R., Besancon, E., Luo, B. H., Stins, M. F., Wang, X., Dedhar, S., & Lo, E. H. (2008). Neuroprotection via matrix-trophic coupling between cerebral endothelial cells and neurons. Proceedings of the National Academy of Sciences of the United States of America. 10.1073/pnas.0801105105

Hernandez-Guillamon, M., Garcia-Bonilla, L., Solé, M., Sosti, V., Parés, M., Campos, M., Ortega-Aznar, A., Domínguez, C., Rubiera, M., Ribó, M., Quintana, M., Molina, C. A., Alvarez-Sabín, J., Rosell, A., Unzeta, M., & Montaner, J. (2010). Plasma VAP-1/SSAO Activity Predicts Intracranial Hemorrhages and Adverse Neurological Outcome After Tissue Plasminogen Activator Treatment in Stroke. Stroke, 41(7), 1528–1535. 10.1161/STROKEAHA.110.584623

Hsieh, H., Boehm, J., Sato, C., Iwatsubo, T., Tomita, T., Sisodia, S., & Malinow, R. (2006). AMPAR removal underlies Abeta-induced synaptic depression and dendritic spine loss. Neuron, 52(5), 831–843. 10.1016/j.neuron.2006.10.035

Hu, X., Ballo, L., Pietila, L., Viesselmann, C., Ballweg, J., Lumbard, D., Stevenson, M., Merriam, E., & Dent, E. W. (2011). BDNF-induced increase of PSD-95 in dendritic spines requires dynamic microtubule invasions. The Journal of Neuroscience: The Official Journal of the Society for Neuroscience, 31(43), 15597–15603. 10.1523/JNEUROSCI.2445-11.2011

Iadecola, C. (2010). The overlap between neurodegenerative and vascular factors in the pathogenesis of dementia. Acta Neuropathologica, 120(3), 287–296. 10.1007/s00401-010-0718-6

Iadecola, C. (2017). The Neurovascular Unit Coming of Age: A Journey through Neurovascular Coupling in Health and Disease. Neuron, 96(1), 17–42. 10.1016/j.neuron.2017.07.030

Jalkanen, S., & Salmi, M. (2008). VAP-1 and CD73, Endothelial Cell Surface Enzymes in Leukocyte Extravasation. Arteriosclerosis, Thrombosis, and Vascular Biology, 28(1), 18–26. 10.1161/ATVBAHA.107.153130

Jin. (2020). Regulation of BDNF-TrkB Signaling and Potential Therapeutic Strategies for Parkinson’s Disease. Journal of Clinical Medicine. 10.3390/jcm9010257

Kadry, H., Noorani, B., & Cucullo, L. (2020). A blood–brain barrier overview on structure, function, impairment, and biomarkers of integrity. Fluids and Barriers of the CNS, 17(1), 69. 10.1186/s12987-020-00230-3

Kellner, Y., Goedecke, N., Dierkes, T., Thieme, N., Zagrebelsky, M., & Korte, M. (2014). The BDNF effects on dendritic spines of mature hippocampal neurons depend on neuronal activity. Frontiers in Synaptic Neuroscience, 6. https://www.frontiersin.org/articles/10.3389/fnsyn.2014.00005

Kim, H., Li, Q., Hempstead, B. L., & Madri, J. A. (2004). Paracrine and Autocrine Functions of Brain-derived Neurotrophic Factor (BDNF) and Nerve Growth Factor (NGF) in Brain-derived Endothelial Cells *. Journal of Biological Chemistry, 279(32), 33538–33546. 10.1074/jbc.M404115200

Kisler, K., Nelson, A. R., Montagne, A., & Zlokovic, B. V. (2017). Cerebral blood flow regulation and neurovascular dysfunction in Alzheimer disease. Nature Reviews Neuroscience, 18(7), 419–434. 10.1038/nrn.2017.48

Kunze, R., & Marti, H. H. (2019). Angioneurins – Key regulators of blood–brain barrier integrity during hypoxic and ischemic brain injury. Progress in Neurobiology, 178. 10.1016/j.pneurobio.2019.03.004

Levine, E. S., Crozier, R. A., Black, I. B., & Plummer, M. R. (1998). Brain-derived neurotrophic factor modulates hippocampal synaptic transmission by increasing N-methyl-D-aspartic acid receptor activity. Proceedings of the National Academy of Sciences of the United States of America, 95(17), 10235–10239. 10.1073/pnas.95.17.10235

Lituma, P. J., Kwon, H.-B., Alviña, K., Luján, R., & Castillo, P. E. (2021). Presynaptic NMDA receptors facilitate short-term plasticity and BDNF release at hippocampal mossy fiber synapses. eLife, 10, e66612. 10.7554/eLife.66612

Lyles, G. A. (1996). Mammalian plasma and tissue-bound semicarba ide-sensitive amine oxidases: Biochemical, pharmacological and toxicological aspects. The International Journal of Biochemistry & Cell Biology, 28(3), 259–274. 10.1016/1357-2725(95)00130-1

Lynch, G., Rex, C. S., Chen, L. Y., & Gall, C. M. (2008). The substrates of memory: Defects, treatments, and enhancement. En European Journal of Pharmacology (Vol. 585, Número 1, p. 2–13). Eur J Pharmacol. 10.1016/j.ejphar.2007.11.082

Ma, Q., Zhao, Z., Sagare, A. P., Wu, Y., Wang, M., Owens, N. C., Verghese, P. B., Herz, J., Holtzman, D. M., & Zlokovic, B. V. (2018). Blood-brain barrier-associated pericytes internalize and clear aggregated amyloid-β42 by LRP1-dependent apolipoprotein E isoform-specific mechanism. Molecular Neurodegeneration, 13(1), 57. 10.1186/s13024-018-0286-0

Malagelada, C., Xifró, X., Miñano, A., Sabriá, J., & Rodríguez-Alvarez, J. (2005). Contribution of caspase-mediated apoptosis to the cell death caused by oxygen– glucose deprivation in cortical cell cultures. Neurobiology of Disease, 20(1), 27–37. 10.1016/j.nbd.2005.01.028

Marie, C., Pedard, M., Quirié, A., Tessier, A., Garnier, P., Totoson, P., & Demougeot, C. (2018). Brain-derived neurotrophic factor secreted by the cerebral endothelium: A new actor of brain function? Journal of Cerebral Blood Flow and Metabolism: Official Journal of the International Society of Cerebral Blood Flow and Metabolism, 38(6), 935–949. 10.1177/0271678X18766772

Martínez, A., Alcántara, S., Borrell, V., Del Río, J. A., Blasi, J., Otal, R., Campos, N., Boronat, A., Barbacid, M., Silos-Santiago, I., & Soriano, E. (1998). TrkB and TrkC signaling are required for maturation and synaptogenesis of hippocampal connections. The Journal of Neuroscience: The Official Journal of the Society for Neuroscience, 18(18), 7336–7350. 10.1523/JNEUROSCI.18-18-07336.1998

Mészáros, Z., Karádi, I., Csányi, A., Szombathy, T., Romics, L., & Magyar, K. (1999). Determination of human serum semicarbazide-sensitive amine oxidase activity: A possible clinical marker of atherosclerosis. European Journal of Drug Metabolism and Pharmacokinetics, 24(4), 299–302. 10.1007/BF03190036

Minichiello, L. (2009). TrkB signalling pathways in LTP and learning. Nature Reviews Neuroscience, 10(12), Article 12. 10.1038/nrn2738

Monnier, A., Prigent-Tessier, A., Quirié, A., Bertrand, N., Savary, S., Gondcaille, C., Garnier, P., Demougeot, C., & Marie, C. (2017). Brain-derived neurotrophic factor of the cerebral microvasculature: A forgotten and nitric oxide-dependent contributor of brain-derived neurotrophic factor in the brain. Acta Physiologica. 10.1111/apha.12743

Noe, C. R., Noe-Letschnig, M., Handschuh, P., Noe, C. A., & Lanzenberger, R. (2020). Dysfunction of the Blood-Brain Barrier—A Key Step in Neurodegeneration and Dementia. Frontiers in Aging Neuroscience, 12. https://www.frontiersin.org/articles/10.3389/fnagi.2020.00185

Novkovic, T., Mittmann, T., & Manahan-Vaughan, D. (2015). BDNF contributes to the facilitation of hippocampal synaptic plasticity and learning enabled by environmental enrichment. Hippocampus, 25(1), 1–15. 10.1002/hipo.22342

Ogura, Y., Sato, K., Kawashima, K.-I., Kobayashi, N., Imura, S., Fujino, K., Kawaguchi, H., & Nedachi, T. (2014). Subtoxic levels of hydrogen peroxide induce brain-derived neurotrophic factor expression to protect PC12 cells. BMC Research Notes, 7, 840. 10.1186/1756-0500-7-840

Pannecoeck, R., Serruys, D., Benmeridja, L., Delanghe, J. R., Geel, N. van, Speeckaert, R., & Speeckaert, M. M. (2015). Vascular adhesion protein-1: Role in human pathology and application as a biomarker. Critical Reviews in Clinical Laboratory Sciences, 52(6), 284–300. 10.3109/10408363.2015.1050714

Parsons, M. P., Kang, R., Buren, C., Dau, A., Southwell, A. L., Doty, C. N., Sanders, S. S., Hayden, M. R., & Raymond, L. A. (2014). Bidirectional Control of Postsynaptic Density-95 (PSD-95) Clustering by Huntingtin *. Journal of Biological Chemistry, 289(6), 3518–3528. 10.1074/jbc.M113.513945

Rauskolb, S., Zagrebelsky, M., Dreznjak, A., Deogracias, R., Matsumoto, T., Wiese, S., Erne, B., Sendtner, M., Schaeren-Wiemers, N., Korte, M., & Barde, Y.-A. (2010). Global deprivation of brain-derived neurotrophic factor in the CNS reveals an area-specific requirement for dendritic growth. The Journal of Neuroscience: The Official Journal of the Society for Neuroscience, 30(5), 1739–1749. 10.1523/JNEUROSCI.5100-09.2010

Robinet, C., & Pellerin, L. (2011). Brain-derived neurotrophic factor enhances the hippocampal expression of key postsynaptic proteins in vivo including the monocarboxylate transporter MCT2. Neuroscience, 192, 155–163. 10.1016/j.neuroscience.2011.06.059

Sagare, A. P., Bell, R. D., & Zlokovic, B. V. (2013). Neurovascular Defects and Faulty Amyloid-β Vascular Clearance in Alzheimer’s Disease. Journal of Alzheimer’s Disease, 33(s1), S87–S100. 10.3233/JAD-2012-129037

Scheffer, S., Hermkens, D. M. A., Van Der Weerd, L., De Vries, H. E., & Daemen, M. J. A. P. (2021). Vascular Hypothesis of Alzheimer Disease: Topical Review of Mouse Models. Arteriosclerosis, thrombosis, and vascular biology, 41(4), 1265–1283. 10.1161/ATVBAHA.120.311911

Schultz, N., Brännström, K., Byman, E., Moussaud, S., Nielsen, H. M., Bank, T. N. B., Olofsson, A., & Wennström, M. (2018). Amyloid-beta 1-40 is associated with alterations in NG2+ pericyte population ex vivo and in vitro. Aging Cell, 17(3), e12728. 10.1111/acel.12728

Shankar, G. M., Li, S., Mehta, T. H., Garcia-Munoz, A., Shepardson, N. E., Smith, I., Brett, F. M., Farrell, M. A., Rowan, M. J., Lemere, C. A., Regan, C. M., Walsh, D. M., Sabatini, B. L., & Selkoe, D. J. (2008). Amyloid-beta protein dimers isolated directly from Alzheimer’s brains impair synaptic plasticity and memory. Nat Med, 14(8), 837–842. 10.1038/nm1782

Smith, D. J., Salmi, M., Bono, P., Hellman, J., Leu, T., & Jalkanen, S. (1998). Cloning of Vascular Adhesion Protein 1 Reveals a Novel Multifunctional Adhesion Molecule. Journal of Experimental Medicine, 188(1), 17–27. 10.1084/jem.188.1.17

Solé, M., Esteban-Lopez, M., Taltavull, B., Fábregas, C., Fadó, R., Casals, N., Rodríguez-Álvarez, J., Miñano-Molina, A. J., & Unzeta, M. (2019). Blood-brain barrier dysfunction underlying Alzheimer’s disease is induced by an SSAO/VAP-1-dependent cerebrovascular activation with enhanced Aβ deposition. Biochimica et Biophysica Acta - Molecular Basis of Disease, 1865(9), 2189–2202. 10.1016/j.bbadis.2019.04.016

Solé, M., Hernandez-Guillamon, M., Boada, M., & Unzeta, M. (2008). P53 phosphorylation is involved in vascular cell death induced by the catalytic activity of membrane-bound SSAO/VAP-1. Biochimica et Biophysica Acta (BBA) - Molecular Cell Research, 1783(6), 1085–1094. 10.1016/j.bbamcr.2008.02.014

Solé, M., Miñano-Molina, A. J., & Unzeta, M. (2015). A cross-talk between Aβ and endothelial SSAO/VAP-1 accelerates vascular damage and Aβ aggregation related to CAA-AD. Neurobiology of Aging, 36(2), 762–775. 10.1016/j.neurobiolaging.2014.09.030

Sun, P., Esteban, G., Inokuchi, T., Marco-Contelles, J., Weksler, B. B., Romero, I. A., Couraud, P. O., Unzeta, M., & Solé, M. (2015). Protective effect of the multitarget compound DPH-4 on human SSAO/VAP-1-expressing hCMEC/D3 cells under oxygen-glucose deprivation conditions: An in vitro experimental model of cerebral ischaemia. British Journal of Pharmacology, 172(22), 5390–5402. 10.1111/bph.13328

Sweeney, M. D., Sagare, A. P., & Zlokovic, B. V. (2018). Blood–brain barrier breakdown in Alzheimer disease and other neurodegenerative disorders. Nature Reviews Neurology, 14(3), Article 3. 10.1038/nrneurol.2017.188

Takei, N., Numakawa, T., Kozaki, S., Sakai, N., Endo, Y., Takahashi, M., & Hatanaka, H. (1998). Brain-derived neurotrophic factor induces rapid and transient release of glutamate through the non-exocytotic pathway from cortical neurons. The Journal of Biological Chemistry, 273(42), 27620–27624. 10.1074/jbc.273.42.27620

Tartaglia, N., Du, J., Tyler, W. J., Neale, E., Pozzo-Miller, L., & Lu, B. (2001). Protein Synthesis-dependent and -independent Regulation of Hippocampal Synapses by Brain-derived Neurotrophic Factor *. Journal of Biological Chemistry, 276(40), 37585–37593. 10.1074/jbc.M101683200

Unzeta, M., Hernàndez-guillamon, M., Sun, P., & Solé, M. (2021). SSAO / VAP-1 in Cerebrovascular Disorders: A Potential Therapeutic Target for Stroke and Alzheimer’s Disease. International Journal of Molecular Sciences, 22(7):3365. 10.3390/ijms22073365

Valente, T., Gella, A., Solé, M., Durany, N., & Unzeta, M. (2012). Immunohistochemical study of semicarbazide-sensitive amine oxidase/vascular adhesion protein-1 in the hippocampal vasculature: Pathological synergy of Alzheimer’s disease and diabetes mellitus. Journal of Neuroscience Research, 90(10), 1989–1996. 10.1002/jnr.23092

Vicario-Abejón, C., Collin, C., McKay, R. D., & Segal, M. (1998). Neurotrophins induce formation of functional excitatory and inhibitory synapses between cultured hippocampal neurons. The Journal of Neuroscience: The Official Journal of the Society for Neuroscience, 18(18), 7256–7271. 10.1523/JNEUROSCI.18-18-07256.1998

von Bohlen und Halbach, O., & von Bohlen und Halbach, V. (2018). BDNF effects on dendritic spine morphology and hippocampal function. Cell and Tissue Research (Vol. 373, Número 3, p. 729–741). Springer Verlag. 10.1007/s00441-017-2782-x

Weksler, B. B., Subileau, E. A., Perrière, N., Charneau, P., Holloway, K., Leveque, M., Tricoire-Leignel, H., Nicotra, A., Bourdoulous, S., Turowski, P., Male, D. K., Roux, F., Greenwood, J., Romero, I. A., & Couraud, P. O. (2005). Blood-brain barrier-specific properties of a human adult brain endothelial cell line. FASEB Journal, 19(13), 1872–1874. 10.1096/fj.04-3458fje

Weksler, B., Romero, I. A., & Couraud, P. O. (2013). The hCMEC/D3 cell line as a model of the human blood brain barrier. Fluids and Barriers of the CNS, 10(1), 1–10. 10.1186/2045-8118-10-16

Yoshii, A., & Constantine-Paton, M. (2014). Postsynaptic localization of PSD-95 is regulated by all three pathways downstream of TrkB signaling. Frontiers in Synaptic Neuroscience, 6. https://www.frontiersin.org/articles/10.3389/fnsyn.2014.00006

Zacchigna, S., Lambrechts, D., & Carmeliet, P. (2008). Neurovascular signalling defects in neurodegeneration. Nat Rev Neurosci, 9(3), 169–181. 10.1038/nrn2336

Zagrean, A.-M., Hermann, D. M., Opris, I., Zagrean, L., & Popa-Wagner, A. (2018). Multicellular Crosstalk Between Exosomes and the Neurovascular Unit After Cerebral Ischemia. Therapeutic Implications. Frontiers in Neuroscience, 12, 811. 10.3389/fnins.2018.00811

Zagrebelsky, M., & Korte, M. (2014). Form follows function: BDNF and its involvement in sculpting the function and structure of synapses. Neuropharmacology, 76, 628–638. 10.1016/j.neuropharm.2013.05.029

Zakharenko, S. S., Patterson, S. L., Dragatsis, I., Zeitlin, S. O., Siegelbaum, S. A., Kandel, E. R., & Morozov, A. (2003). Presynaptic BDNF required for a presynaptic but not postsynaptic component of LTP at hippocampal CA1-CA3 synapses. Neuron, 39(6), 975–990. 10.1016/s0896-6273(03)00543-9

Ziegenhorn, A. A., Schulte-Herbrüggen, O., Danker-Hopfe, H., Malbranc, M., Hartung, H. D., Anders, D., Lang, U. E., Steinhagen-Thiessen, E., Schaub, R. T., & Hellweg, R. (2007). Serum neurotrophins—A study on the time course and influencing factors in a large old age sample. Neurobiology of aging, 28(9), 1436–1445. 10.1016/J.NEUROBIOLAGING.2006.06.011

Zlokovic, B. V. (2011). Neurovascular pathways to neurodegeneration in Alzheimer’s disease and other disorders. Nature Reviews Neuroscience, 12(12), Article 12. 10.1038/nrn3114

